# Transcriptional landscape of the archaeal cell cycle is broadly conserved in eukaryotes

**DOI:** 10.1101/2024.11.20.624502

**Authors:** Miguel V. Gomez-Raya-Vilanova, Jérôme Teulière, Sofia Medvedeva, Yuping Dai, Eduardo Corel, Philippe Lopez, François-Joseph Lapointe, Debashish Bhattacharya, Louis-Patrick Haraoui, Elodie Turc, Marc Monot, Virginija Cvirkaite-Krupovic, Eric Bapteste, Mart Krupovic

## Abstract

The cell cycle is a series of events that occur from the moment of cell birth to cell division. In eukaryotes, cell growth, genome replication, genome segregation, and cytokinesis are strictly coordinated, defining discrete cell cycle phases. In contrast, these key processes may occur concurrently in bacteria. Thermoacidophilic archaea in the genus *Saccharolobus* follow a defined cell cycle program, with the first pre-replicative growth (G1) phase, followed by the chromosome replication (S) phase, the second growth (G2) phase, and rapid genome segregation (M) and cytokinesis (D) phases. However, whether other processes, such as metabolism, catabolism, protein translation, and antiviral defense also occur at specific cell cycle phases, as in eukaryotes, or are active throughout the cell cycle, as in bacteria, remains unclear. To address this question, we synchronized cultures of *S. islandicus* and performed an in-depth transcriptomic analysis of samples enriched in cells undergoing the M-G1, S, and G2 phases. Differential gene expression and consensus gene co-expression network analyses provided a holistic view of the *S. islandicus* cell cycle. In addition to the core transcriptome network, which is expressed throughout the cell cycle, we show that diverse metabolic pathways, protein synthesis, cell motility and even antiviral defense systems, are expressed in a cell cycle dependent fashion. Our data also refines understanding of the processes previously known to be linked to the cell cycle, such as DNA replication. We show that most DNA replication genes are expressed prior to the S phase, during the M-G1, whereas expression of the major chromatin genes, and accordingly, chromatinization are concomitant with replication. A statistical model was used to define sets of signature genes characteristic of each of the analyzed cell cycle phases, emphasizing transcriptional stratification of the phases. Signature genes are more conserved across Thermoproteota than non-signature genes and their peak expression, especially for the M-G1 and G2 specific genes, matches that of homologs in yeast. Collectively, our data elucidate the complexity of the *S. islandicus* cell cycle and suggest that it more closely resembles the cell cycle of eukaryotes than previously appreciated.

## INTRODUCTION

The life of a cell unfolds through a series of intricately coordinated events that culminate in the production of two daughter cells. Faithful execution of this program, known as the cell cycle, ensures the perpetuation of cellular life. The core processes essential for cell cycle progression, such as accumulation of biomass, genome replication and cytokinesis, are common to all organisms^1–3^, but their coordination and the underlying molecular mechanisms exhibit remarkable diversity. For instance, eukaryotes encode a diverse set of cyclins^1^, the molecular regulators of the cell cycle, for which no homologs exist in archaea and bacteria^2,3^. Understanding the interplay between different cellular processes and the evolution of these relationships in different cellular domains is of fundamental interest and practical importance, because it offers insights into how both unicellular and multicellular life forms have evolved on our planet and may provide new targets for therapies.

Bacteria and eukaryotes rely on distinct genome replication and cell division machineries, with bacteria using the FtsZ-based system for division^4^ and eukaryotes employing the ESCRT (endosomal sorting complexes required for transport) machinery for membrane abscission during cytokinesis^5,6^. In eukaryotes, genome replication and cytokinesis are typically separated in time with the cell cycle being divided into four phases^1,7,8^: (i) during the first gap (G1) phase the cell grows and prepares for genome replication; (ii) the synthesis of genomic DNA takes place during the S phase; (iii) the second gap (G2) phase is a period of rapid cell growth and protein synthesis; and, finally, (iv) during the mitosis (M) phase the sister chromatids are segregated and the cell is divided in two. Progression through the cell cycle phases is tightly controlled at several checkpoints^1,7,8^, with errors at any of the checkpoints leading to devastating consequences at both cellular and organismal levels, including cell death, cancers and various other pathologies^9–11^.

In bacteria, the cell cycle is traditionally divided into three periods^2,12^: (i) the birth (B) period defined as the time between cell birth and initiation of genome replication, (ii) the C period – from chromosome replication initiation to termination, and (iii) the D period, between completion of DNA replication and cell division. Unlike in eukaryotes, the periods of the bacterial cell cycle are typically less strictly separated in time. For instance, bacterial genome replication is concomitant with segregation of chromosomal DNA into developing daughter cells and, depending on the growth conditions, generation time can be shorter than the combined duration of the C and D periods, leading to multiple DNA replication initiation events and overlapping replication cycles in each cell^13^. Accordingly, bacteria generally lack the cell cycle checkpoints, although in some bacterial models, cell volume and other characteristics (e.g., motility or lack thereof) are important for cell cycle progression^14–16^.

Archaea, single-celled organisms comprising the third cellular domain^17^, display a remarkable diversity of metabolic capabilities, environmental adaptations and molecular machineries responsible for key cellular processes. Similar to bacteria, archaea have circular chromosomes with most genes organized into operons^18^. However, archaeal proteins involved in replication, transcription and translation are more closely related to homologs in eukaryotes^19,20^. By contrast, cell division systems can be either bacterial-like, based on the FtsZ rings^21–23^, or eukaryotic-like, centered around the ESCRT complex^24–27^. Whether archaea display a bacterial-like or eukaryotic-like cell cycle remains unclear due to the scarcity of tractable model systems in this domain of life^18^.

Thermoacidophilic archaea (optimal growth at ∼80°C and pH∼3) in the order Sulfolobales emerged as models for cell biology and cell cycle studies^28^. The coccoid-shaped Sulfolobales cells use the ESCRT machinery for division and follow the eukaryotic-like cell cycle paradigm (Fig. S1A). An exponentially growing Sulfolobales cell starts the cycle with a short (<5% of the cell cycle) pre-replicative G1 phase, which is followed by the genome replication S phase (30-35% of the cycle). Then, the cell enters the longest G2 phase (>50% of the whole cycle) during which the cell prepares for genome segregation. Finally, the cycle culminates with two short, M and D, phases (each lasting <5% of the cycle) during which the genome copies are segregated and the cell is divided^18,28^. The overall program of the cell cycle appears to be conserved throughout the class Thermoprotei (formerly known as phylum Crenarchaeota)^29^. Importantly, the cell cycle in a Sulfolobales population can be synchronized using a transient treatment with acetic acid, which presumably induces respiration uncoupling, arresting the cells in the post-replicative G2 cell cycle phase. Acetic acid removal allows the near-synchronous resumption of the cell cycle^28^. Whereas the overall outline of the Sulfolobales cell cycle and coordination between genome replication and cytokinesis have been defined^18,25,30–33^, it remains unknown whether other central cellular processes, such as diverse metabolic and catabolic pathways or protein translation, are harmonized with the cell cycle.

To obtain a more integrated view of the different processes taking place during the archaeal cell cycle, we performed a deep transcriptomic analysis using *Saccharolobus islandicus* (formerly *Sulfolobus islandicus*; order Sulfolobales) as a model. We analyzed differential gene expression patterns during distinct cell cycle phases and harnessed the power of the consensus Gene Coexpression Networks (GCN), a state-of-the art statistical framework enabling analysis of negative and positive correlations between all expressed genes. The two complementary analytical approaches showed that the cell cycle of *S. islandicus* more closely resembles that of eukaryotes than previously appreciated. We show that not only replication, chromosome segregation and division, but other cellular processes occur in a cell cycle-dependent manner. Finally, we used a machine learning approach to identify signature genes which can be used as markers of specific cell cycle phases. Remarkably, these genes were generally well-conserved across Thermoproteota and their timing of expression matched the peak of expression of their homologs in yeast. Collectively, our data provide a robust understanding of the *S. islandicus* cell cycle, opening new avenues for future research.

## RESULTS AND DISCUSSION

### Overview of the transcriptional landscape across cell cycle phases

To analyze how *S. islandicus* cells coordinate different processes as they progress through the cell cycle, we sequenced the transcriptomes at three time points during which the populations were enriched in cells undergoing the M-G1, S and G2 phases, as evidenced by flow cytometry (Supplementary Fig. S1, Supplementary Fig. S2A). Due to the short duration of the consecutive M, D and G1 phases, populations specifically enriched in these phases could not be collected separately. Hence, the corresponding populations were pooled together within a single sample, denoted herein as M-G1. To obtain additional information about the coordination of gene expression, we leveraged the statistical framework of the consensus Gene Coexpression Networks (GCN). Out of the total of 2630 predicted genes, including protein coding genes, tRNA, rRNA and ncRNA, 2558 were expressed (i.e., at least one read per million reads), with 2356 genes being expressed in all three phases (Supplementary Fig. S2B, Supplementary table S1). Principal component analysis showed that 56.4% of the variance can be explained by the three first principal components and that samples can be clustered according to the cell cycle phases (Supplementary Fig. S2C).

The *S. islandicus* chromosome is structurally organized into A and B compartments that have high and low gene expression, respectively^34,35^. Consistently, genes in the A compartment had on average higher expression than genes in the B compartment during all cell cycle phases (Supplementary Fig. S2D), with the expression being the highest near the three origins of replication (*ori*). However, there was only negligible difference (t-test, *p*=0.07) in the overall level of expression within the *ori* proximal regions during different cell cycle phases.

Pairwise comparison of the gene expression profiles during consecutive cell cycle phases, namely, M-G1 vs S, S vs G2 and G2 vs M-G1, revealed 743 differentially expressed genes, which were either up- or down-regulated in a particular cell cycle phase (Fig. 1A; Supplementary Fig. S3), suggesting that expression of 28% of *S. islandicus* genes follows a coordinated pattern throughout the cell cycle. Of the 743 differentially expressed genes, 308 were specifically up- or down-regulated during a particular cell cycle phase, pointing to marked differences between the processes that occur during the three phases. The M-G1 phase displayed the highest number of differentially expressed genes when compared to either S or G2 phase (n=198 in M-G1 vs n=3 in S and n=107 in G2).

**Figure 1.**
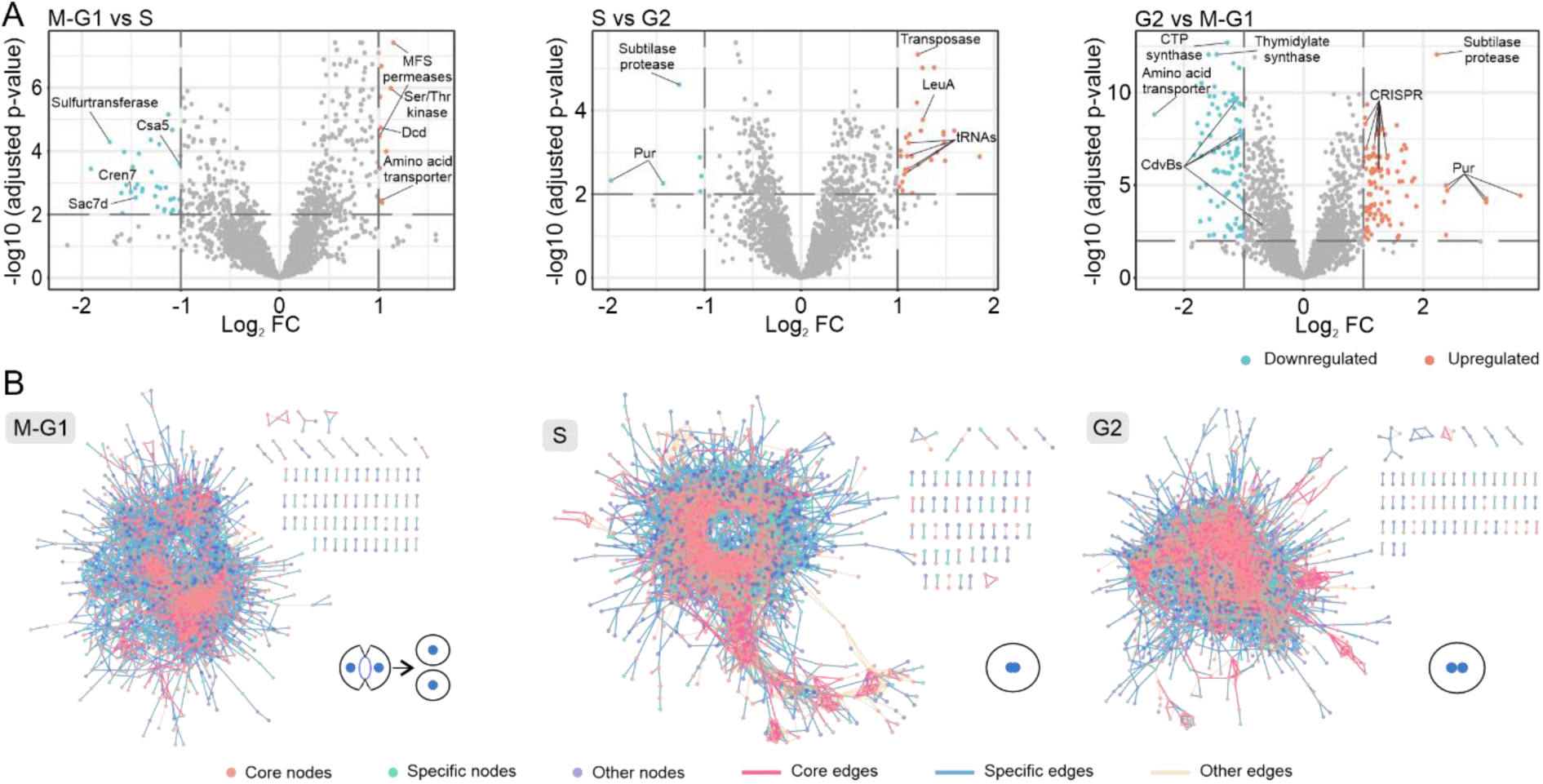
Overview of the transcriptional landscape across cell cycle phases. **A.** Differential gene expression by pair-wise comparison of the three analyzed phases (M-G1, S and G2) represented using volcano plots. The y-axis represents the adjusted *p* value in logarithmic scale in base 10. The x-axis represents the fold change (FC) between the two phases in logarithmic scale in base 2. The horizontal lines mark the thresholds for significance, i.e., a *p* value of <1 (-log_10_(0.01) = 2). The vertical lines mark the thresholds for strong differential expression, i.e., FC of 2 (log_2_(±2) = ±1). Genes of interest are marked with general annotations. **B.** Consensus gene co-expression networks of the three analyzed phases (M-G1, S and G2). Gene co-expression networks constructed for each of the three phases consist of nodes (genes) connected by edges. An edge will be drawn when two nodes display significant co-expression. Nodes and edges are classified according to their presence in one (specific), two (other) or all the networks (core). The presumed morphological states characterizing the different cell cycle phases are shown next to the corresponding networks.

Differential gene expression (DGE) analysis provides information on the changes in the extent of expression of genes during a given cell cycle phase. However, although instructive, this information does not capture the subtle changes in the rewiring of gene co-expression patterns, which might have a major impact on the progression of the cell cycle. Thus, to gain an orthogonal view on the co-expression of genes during different phases of the cell cycle, we constructed consensus GCNs for each of the three phases. Each GCN consists of nodes and edges, where nodes are genes and edges, i.e., lines connecting the nodes, represent statistically significant positive or negative correlations between the expression values of the connected genes. Each GCN had a different number of nodes and edges, which were classified into (i) the ‘core’, i.e., common to all three phases, (ii) ‘phase-specific’, i.e., co-expressed during one of the phases or (iii) ‘other’, if the co-expression was detected during two of the three phases (Fig. 1B). This analysis revealed that the S phase is characterized by the most complex network containing the highest number of edges and nodes (n=25,100 and n=2,073, respectively; Supplementary table S1), indicative of higher coordination of gene expression during this phase compared to the other phases. The DGE and GCN analyses show that a considerable fraction of genes follows cell cycle-dependent patterns of expression that manifest either at the level of expression strength or co-expression wiring.

### Composition and properties of the core network

Contrasting the phase-dependent expression of genes that, by definition, characterize a particular cell cycle phase, the genes and groups of genes that ensure the maintenance of cell viability are expected to be co-expressed throughout the cell cycle. This co-expression is expected to occur regardless of whether the products of the co-expressed genes are functionally coupled, i.e., function in the same process or pathway. Analysis of such constitutively co-expressed genes, which we refer to as the housekeeping ‘core’ network, allowed us to define the house-keeping tasks necessary for the functioning of a cell. The core GCN, consisting of the core nodes connected by the core edges, contained 417 genes (Supplementary table S2; note that only groups of four or more co-expressing genes were included as part of the core network). The core network included 1,255 core edges, so that each core gene on average was co-expressed with 3 other core genes. Analysis of the overall expression levels of the genes forming the core network revealed notable variation in expression levels across cell cycle phases, with significantly higher activity during the S and G2 phases (*p* value < 0.0001; Fig. 2A).

**Figure 2.**
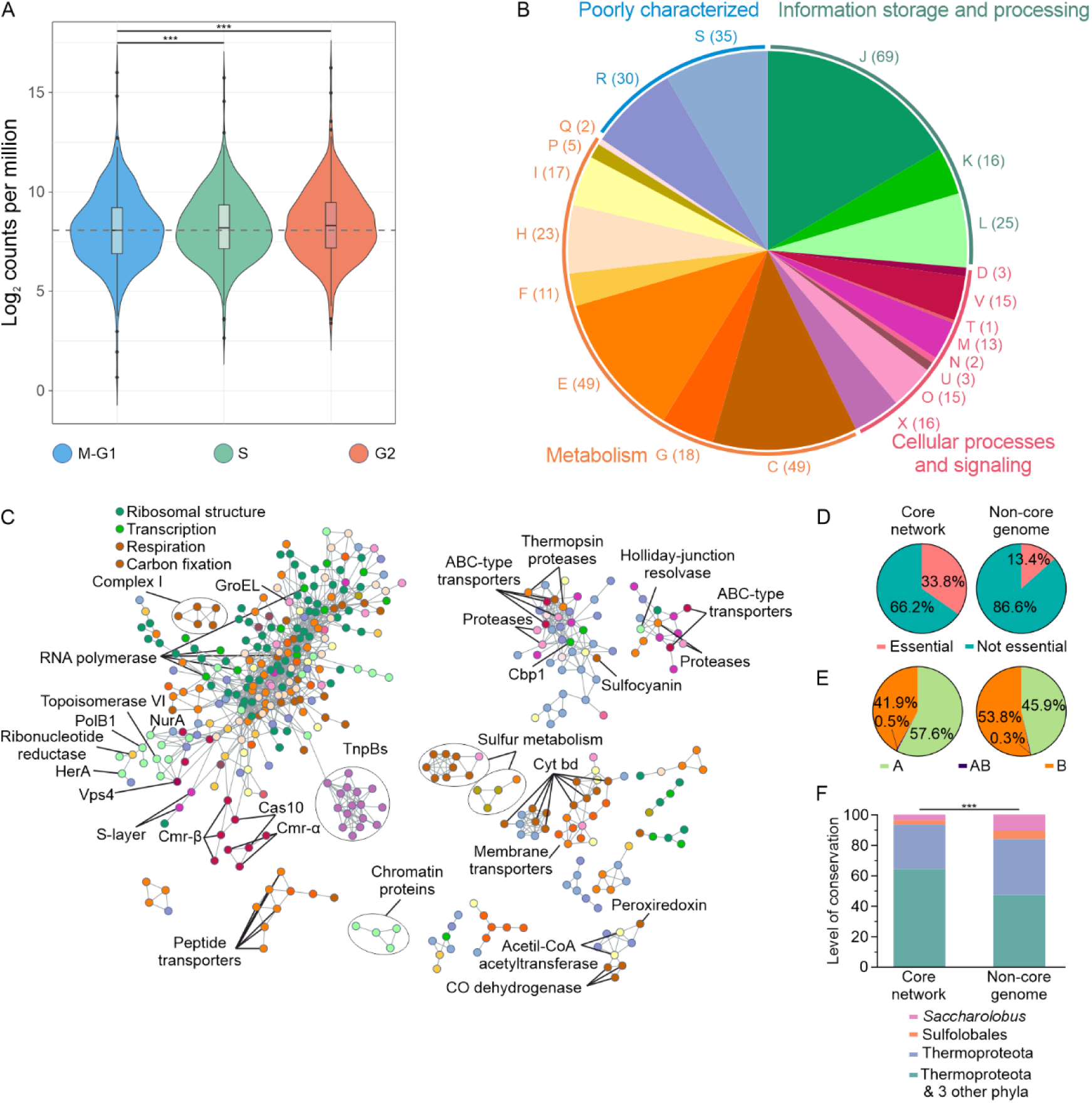
Core network of *S. islandicus*. **A.** Expression of the genes included in the core network throughout the three analyzed phases (M-G1, S and G2). Expression is represented by violin plots in which the y-axis represents the expression in counts per million reads in logarithmic scale in base 2. A dashed horizontal line marks the average of the M-G1 phase as a reference. Statistical significance was calculated by performing a Wilcoxon test *** = *p* value <0.0001 **B.** Functional classification of the core genes. Genes were annotated and classified using the arCOG framework (see Methods). A pie chart shows the number of genes assigned to each of the 26 arCOG categories. A correspondence between the letter code and the name of each category can be found in Supplementary table S1. The different categories are grouped into four classes: (i) Information storage and processing, (ii) Cellular processes and signaling, (iii) metabolism and (iv) poorly characterized. **C.** Topological representation of the core network. Each node is colored according to its arCOG category following the same color coding as in panel B. Genes of interest are marked. **D.** Essentiality of the genes in the core network compared to the non-core genes. Two pie charts show the percentage of essential genes in the core and non-core genomes. **E.** Compartmentalization of the genes in the core network compared to the non-core genes. Information on compartmentalization of the genome was extracted from data obtained previously^35^. Two pie charts show the percentage of genes in each of the two compartments in the core and non-core genomes. Unassigned genes are classified as AB. **F.** Conservation of the genes in the core network compared to the non-core genes. Genes were assigned to one of the four categories according to their conservation: (i) genes exclusive to the genus *Saccharolobus*, (ii) those restricted to the order Sulfolobales, (iii) genes conserved across the phylum Thermoproteota, and (iv) genes present in Thermoproteota and three other archaeal phyla, namely, Methanobacteriota, Halobacteriota and Thermoplasmatota. Two bar plots show the percentage of genes in each of the four categories for the core and non-core genomes. Statistical significance was calculated by performing a Chi-square test comparing the conservation distribution of the non-core genome with the core genes. *** = *p* value <0.0001

All the nodes (i.e., genes) in the core network were assigned to functional categories according to the archaeal Clusters of Orthologous Genes (arCOG) annotation^36^ (Fig. 2B). Only ∼15% of the genes in the core network could not be functionally annotated (arCOG categories R and S). Most genes in the core network were associated with central cellular processes such as translation, replication, energy production and various metabolic pathways. Analysis of the network topology, revealed several differentiated subclusters (Fig. 2C), suggesting the existence of transcriptional finetuning of functionally coupled genes. For instance, the largest subcluster was enriched in genes responsible for ribosome structure (e.g., ribosomal proteins uL23, uL2, uS3, uS8), transcription (e.g., transcription elongation factor NusA, ribonuclease PH, DNA-directed RNA polymerase), genome replication and repair (e.g., DNA polymerase B1, topoisomerase VI, HerA, NurB), respiration (e.g., ATP-synthase, succinate dehydrogenase), carbon fixation (e.g., succinyl-CoA synthetase, aconitase A) and included genes responsible for the maintenance of cell envelope (S-layer protein) (Fig. 2C). By contrast, several ABC-type transporters and sulfur metabolism genes (e.g., heterodisulfide reductase, sulfite reductase, ATP-sulfurylase), although also included in the core network, formed separate GCN subclusters. Notably, some of the CRISPR defense system genes (e.g., Cas10, Cmr6g7, Cmr1g7 and Cmr5SS of the Cmr-β cassette) were part of the core network, included in the largest subcluster, whereas other *cas* genes were expressed during particular cell cycle phases (see below), suggesting subtle transcriptional control of this complex defense system.

We assessed whether the core network is enriched in essential genes. To this end, we took advantage of the genome-wide gene essentiality information available for a closely related *S. islandicus* strain M.16.4^37^ and compared the fraction of essential genes in the core network to that of the non-core genes. The proportion of essential genes was more than twice higher in the core network than in the rest of the genome (34% vs 13%, *p* value < 0.01; Fig. 2D). The majority (n=115, 82%) of the essential genes were found in the largest subcluster of co-expressed core genes (Supplementary Fig. S4A), suggesting the existence of a mechanism ensuring coordinated and stable co-expression of genes that are critical for the cell functions. We assessed the distribution of the core genes with respect to the chromosomal A and B compartments. Slightly more than half (57.6%) of the core network genes were localized in the A compartment (*p* value < 0.01; Fig. 2E). Collectively, these observations suggest that the *S. islandicus* chromosome has evolved to accommodate essential genes important throughout the cell cycle in the transcriptionally active, less tightly condensed part of the chromosome.

We then evaluated to what extent the core network is conserved in other archaea. To this end, genes from the core network were assigned to one of the four categories: (i) genes exclusive to the genus *Saccharolobus*, (ii) those restricted to the order Sulfolobales, (iii) genes conserved across the phylum Thermoproteota, and (iv) genes present in Thermoproteota and three other archaeal phyla (as defined by GTDB), namely, Methanobacteriota, Halobacteriota and Thermoplasmatota. We found that the core genes displayed significantly higher conservation compared to non-core genes, with 93.5% of the core genes being conserved across Thermoproteota, with 62.5% of the core genes being also conserved in three other phyla. However, when the non-core genes were considered, the fraction decreased to 83.7% and 46.1%, respectively (*p* value < 0.01; Fig. 2F). Moreover, analysis of the network topology showed that the most widely conserved genes occupied the central position of the largest co-expression subcluster (Supplementary Fig. S4C). Indeed, the degree of the most widely conserved genes was significantly higher than that of less conserved genes (Supplementary Fig. S4D), suggesting that the interplay between the core genes evolved prior to the radiation of archaeal diversity and that new, taxon-specific genes and their interactions with the conserved components of the core network have been established subsequently, likely concomitant with archaeal diversification.

### Many key cellular processes are coordinated with the cell cycle progression in *S. islandicus*

To determine how *S. islandicus* coordinates different processes along the cell cycle, we assessed the differential expression and co-expression during different cell cycle phases of all 2558 expressed genes (Figs. 3A, B and Supplementary Fig. S5A). Additionally, KEGG enrichment analysis facilitated the identification of pathways significantly upregulated during each phase (Fig. 3C, Supplementary Fig. S5B-D).

**Figure 3.**
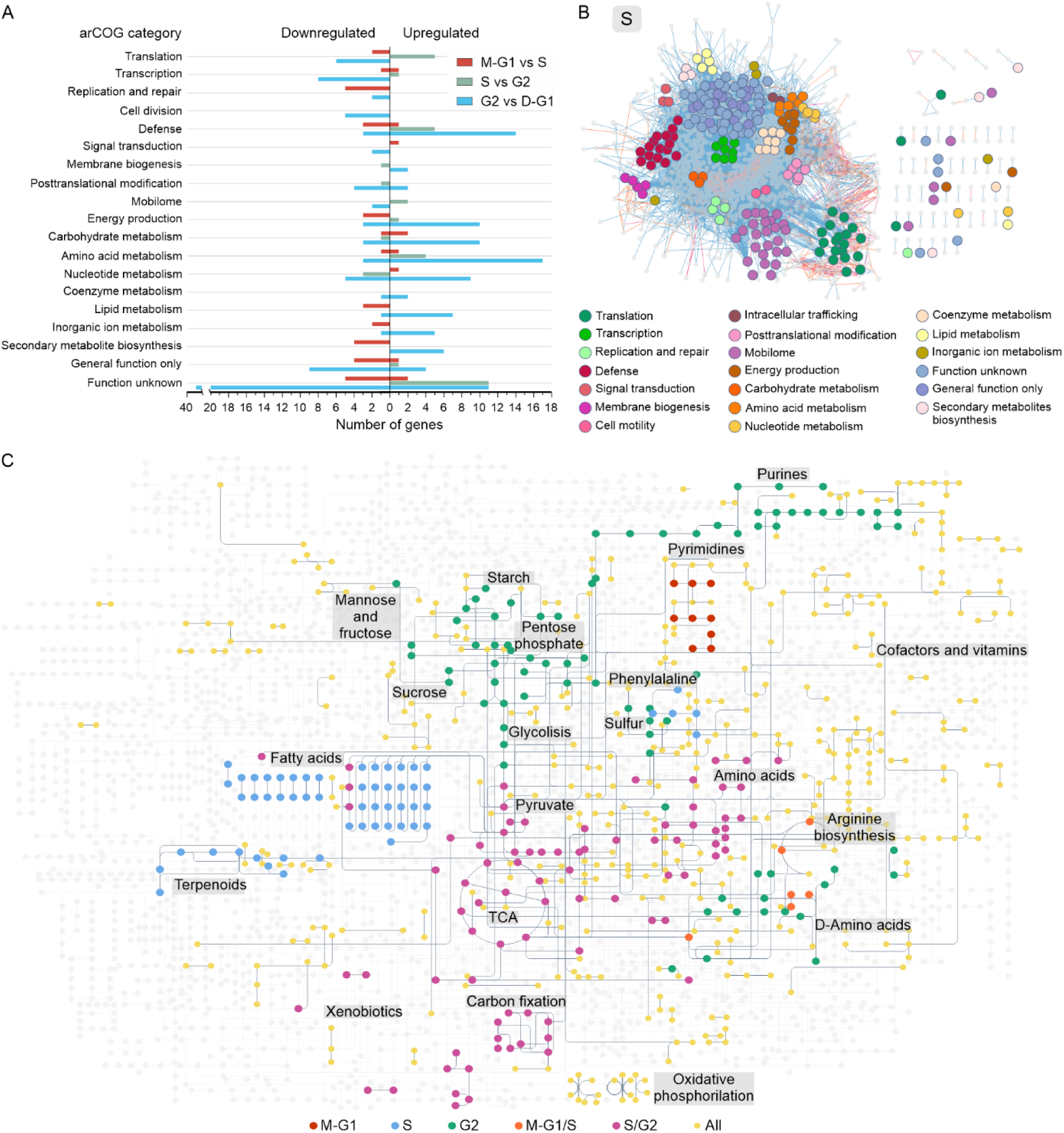
Cell cycle phase specific processes. **A.** Classification of the strongly differentially expressed genes in each pair-wise comparison by arCOG category. Genes were annotated and classified using the arCOG framework (see Methods). Genes were considered strongly differentially expressed when their fold change (FC) was at least ±2 and their adjusted *p* value was <0.01. The number of genes is shown on the x-axis, genes to the right of the vertical axis are upregulated in the corresponding comparison, while genes to the left are downregulated. **B.** Specific co-expression during the S phase. GCN of the S phase with the specific nodes highlighted, grouped and colored by arCOG category (see Supplementary figure S5 for GCNs of the M-G1 and G2 phases). **C.** Differences in the metabolism between phases. A full metabolic map of *S. islandicus* was extracted from the KEGG database^89^ (accession number: sir01100). Each dot represents a metabolite while each line represents the enzymes that transform those metabolites. Each pathway is classified into six groups depending on their peak of activity according to the KEGG enrichment and DGE analysis: (i) ‘All’, if their activity does not change, (ii) ‘M-G1’, (iii) ‘S’ or (iv) ‘G2’, if their activity peaks at each of the corresponding phases, and (v) M-G1/S or (vi) S/G2, if their activity increased at the M-G1 or S phases and was maintained high into the following phase. Different pathways of interest are labeled on the map next to their metabolites.

#### Lipid biosynthesis and membrane biogenesis

Following cell division, the daughter cells start accumulating biomass and increase in size, up until the next round of cell division^18^. Increase in the surface area of the cell necessitates the synthesis of additional lipids. Similar to other members of the Sulfolobales, the *S. islandicus* cell membrane primarily consists of different species of glycerol dibiphytanyl glycerol tetraethers (GDGTs), which contain diverse polar head groups and a variable number of cyclopentane rings in the hydrophobic isoprenoid core^38^. Nine enzymes that participate in the synthesis of GDGT lipids have been identified^39–42^ (Fig. 4A) and five of them are significantly upregulated during the M-G1 phase compared to either S or G2 phase (Fig. 4B). Among these, digeranylgeranylglyceryl phosphate (DGGGP) synthase, which catalyzes the formation of an ether bond linking the second isoprenoid chain to the lipid precursor^43^, and calditol synthase (Cds), responsible for the synthesis of a unique cyclopentyl head group which plays a key role in the acid resistance of Sulfolobales^42^, are the most strongly upregulated genes. Notably, the GCN analysis showed that the lipid biosynthesis pathway is highly coordinated during the M-G1 and S phases (Fig. 4C, inset) and synchronized with expression of diverse membrane proteins, post-translational modification enzymes and various systems embedded in the membrane, such as the Complex II of the electron transport chain and the ATP synthase (Fig. 4C). This observation suggests the existence of a link between cell membrane dependent systems and lipid biosynthesis. During M-G1, we observed higher specific co-expression of genes encoding glycosyltransferases (SiRe_RS03975, SiRe_RS04230, SiRe_RS08195 and SiRe_RS02080), which according to their arCOG annotation are predicted to participate in membrane biogenesis (Supplementary Fig. S5A). Furthermore, many of the genes co-expressed with the lipid biosynthesis enzymes encode poorly characterized proteins, which could represent missing players in the membrane biogenesis processes. For instance, glycosyltransferase (SiRe_RS06775) and N-acetylneuraminate lyase (SiRe_RS10400) identified in the GCN analysis could participate in the synthesis of the lipid headgroups (Fig. 4C).

**Figure 4.**
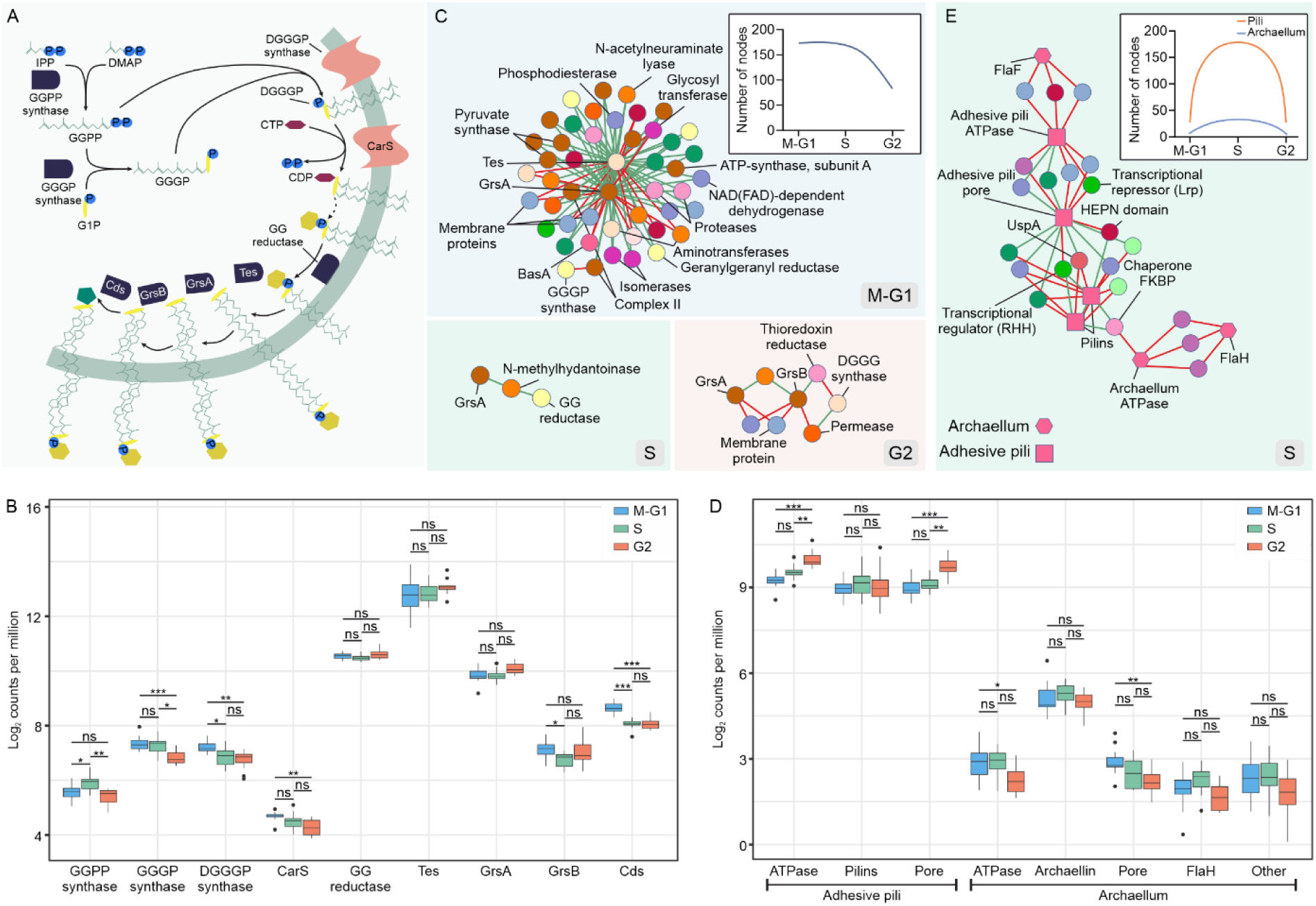
Changes in membrane biogenesis, motility and adhesion across the cell cycle. **A.** Graphical representation of the major known steps of lipid biogenesis in *S. islandicus*. Briefly, isopentenyl pyrophosphate (IPP) or dimethylallyl monophosphate (DMAP) are used by the geranylgeranyl diphosphate (GGPP) synthase to synthetize GGPP. Geranylgeranylglyceryl phosphate (GGGP) is then synthetized by the GGGP synthase by addition of glycerol-1-phosphate. A molecule of GGPP and one of GGGP are used to build digeranylgeranylglyceryl phosphate (DGGGP) by the DGGGP synthase. The polar head of the lipid is then activated by the CDP-diglycerade synthase. After several possible modifications that can happen in the polar head, the terpenoid chain is reduced by the geranylgeranyl (GG) reductase to form archaeol. Two archaeol molecules are transformed by the tetraether synthase (Tes) into a GDGT-0 (zero cyclopentane rings) molecule. Different number of cyclopentane rings can be added to the molecule by GrsA (up to GDGT-4) and by GrsB (from GDGT-4 to GDGT-8). The internal polar head can be modified by the Calditol synthase to form a calditol-linked GDGT. **B.** Expression of the major enzymes participating in lipid biogenesis throughout the cell cycle. Boxplot of the expression of the different enzymes in the 15 replicates. The x-axis represents the counts per million reads in logarithmic scale in base 2. Statistical significance is extracted from the DGE analysis, where it was calculated with the limma package and *p* values were adjusted for multiple comparisons using the Benjamini-Hochberg method. * = *p* value ≤ 0.01. ** = *p* value ≤ 0.001. *** = *p* value ≤ 0.0001. **C.** Gene co-expression subnetworks of the major enzymes participating in membrane biogenesis. A subnetwork is extracted from the full GCN by maintaining only those nodes that co-express with at least two enzymes participating in membrane biogenesis. Nodes are colored by arCOG category following the same color code as in Fig. 2B, while edges are colored green if the co-expression is positive and red if it is negative. **C inset.** Size of the co-expression subnetwork of membrane biogenesis throughout the cell cycle. Line plot representing the number of nodes of the subnetwork of all genes co-expressing with the major membrane biogenesis enzymes at each of the cell cycle phases. **D.** Expression of the adhesive pili and archaellum components. Boxplot of the expression of the different components of the adhesive pili and the archaellum machinery in the 15 replicates. The x-axis represents the counts per million reads in logarithmic scale in base 2. Statistical significance is extracted from the DGE, like in panel B. **E.** Gene co-expression subnetwork at the S phase of the components of the adhesive pili and archaellum. The subnetwork is extracted from the full S GCN by maintaining only those nodes that co-express with at least two components of the adhesive pili or the archaellum. **E inset.** Size of the co-expression subnetwork of motility and adhesion components throughout the cell cycle. Line plots represent the number of nodes of the subnetworks of all genes co-expressing the adhesive pili and the archaellum components at each of the cell cycle phases.

#### Genome replication and architecture

Genome replication is one of the focal points of the cell cycle and, by definition, occurs during the S phase, as evidenced by flow cytometry analysis (Supplementary Fig. S1B). Unexpectedly, some of the key components of the replisome, such as replicative DNA polymerase PolB1 (SiRe_RS07370), replicative minichromosome maintenance (MCM) helicase (SiRe_RS06220), Gins23 (SiRe_RS06225), which participates in replication initiation and elongation, PolB1 binding protein 2 (SiRe_RS07230), and PCNA sliding clamp (SiRe_RS08085), are upregulated during the M-G1, a period preceding the actual S phase (Supplementary table S2). Moreover, Orc1-1 (SiRe_RS08850), Orc1-3 (SiRe_RS00005) and WhiP (SiRe_RS06120), the three replication initiators of *S. islandicus*, are also upregulated during the M-G1, with expression of Orc1-3 and WhiP being also maintained throughout the S. The peak expression of Orc-1-1 and WhiP was observed during M-G1, whereas Orc1-3 expression peaks during the S phase. By contrast, the ATP-dependent DNA ligase (SiRe_RS09250), which ligates the Okazaki fragments during the lagging strand synthesis^20^, is upregulated during the S phase, indicating that some replisome components have a different temporal expression during the cell cycle. Nevertheless, these observations suggest that the cell prepares for DNA replication in advance, by synthesizing most of the necessary enzymes during the M-G1 phase. Alternatively, the replisome components could also participate in DNA repair, preparing the genome for replication during the S phase. Indeed, some of the DNA repair genes are upregulated during the M-G1 phase as well (see Supplementary text).

Many chromatin proteins, including Cren7 (SiRe_RS05625), two Sul7d family proteins (SiRe_RS03405 and SiRe_RS13370), Sul12a family protein (SiRe_07595) and two Sso7c4 homologs (SiRe_07595 and SiRe_RS09950), were strongly upregulated during the S phase (Fig. 3A, Supplementary table S1), suggesting that chromatinization takes place concomitant with DNA replication. It has been recently suggested that Lrs14 family proteins of Sulfolobales should be considered as chromatin organizing proteins^44^. Our data show that Lrs14 in *S. islandicus* (SiRe_RS09945) displays a similar transcription pattern as the main chromatin proteins, supporting the involvement of Lrs14 in chromatin organization. Given that chromatin proteins are among the most abundant proteins in the cell, chromatinization is likely to necessitate extensive protein translation during the S phase. Indeed, many translation-related genes, including those encoding a subset of ribosomal proteins, tRNAs, glycyl-tRNA synthetase, a subunit of the RNase P and translation initiation factor 6 as well as thermosome responsible for protein folding, were upregulated during the S phase (see Supplementary text for details).

#### Central metabolism

We next assessed whether the central metabolic pathways also displayed differential regulation during the cell cycle phases. To this end, we performed the KEGG enrichment analysis (Supplementary Fig. S5B-D) and analyzed the patterns of differential expression of genes assigned to arCOG categories related to metabolism (Fig. 3A). The two approaches provided congruent and complementary results. Many pathways, including those related to biosynthesis of amino acids and nucleotides (see Supplementary text) as well as carbon metabolism, were not uniformly expressed across different cell cycle phases.

The tricarboxylic acid (TCA) cycle (also known as the Krebs or citric acid cycle) is one of the key energy-generating metabolic pathways of the cell. Through a series of biochemical reactions, TCA releases the energy stored in nutrients through the oxidation of acetyl-CoA derived from carbohydrates, fats, and proteins. We found that the carbon fixation pathways, an assemblage of metabolic pathways terminating in the TCA, are upregulated during the S phase and stay active during G2 (Fig. 3C, Supplementary Fig. S5B-D). These pathways fix carbon through the synthesis of malonyl-CoA, which can then be transformed into succinyl-CoA, a key intermediate in the TCA cycle. The subunits of the acetyl-CoA carboxylase (SiRe_RS01265, SiRe_RS01270, SiRe_RS01275), which synthesizes malonyl-CoA, as well as the methylmalonyl-CoA epimerase (SiRe_RS01085) and mutase (SiRe_RS01080), which catalyze the last step in the transformation to succinyl-CoA, are upregulated during the S phase and even more strongly during G2, compared to M-G1 (supplementary table S1). Alternatively, the succinyl-CoA can enter the autotrophic hydroxybutyrate cycle^45^ and produce acetoacetyl-CoA, which will be broken into two acetyl-CoA molecules. Some enzymes participating in this transformation, namely, succinyl-CoA reductase (SiRe_RS04600), succinate semialdehyde reductase (SiRe_RS07755), 3-hydroxybutyryl-CoA dehydrogenase (SiRe_RS11400) and two acetyl-CoA acetyltransferases (SiRe_RS07425 and SiRe_RS13030), follow expression patterns similar to enzymes producing succinyl-CoA. Moreover, although not significantly enriched in any of the phases, the TCA cycle includes several enzymes, such as succinate dehydrogenase (SiRe_RS00780 and SiRe_RS00785) and the aconitate hydratase (SiRe_RS0595), that are also upregulated during the S and G2 phases. These results indicate that the most central and important pathways for production of energy tend to be less active during M-G1.

Notably, the substrates used for energy production during the S and G2 appear to be different. For instance, fatty acid degradation pathways (Supplementary Fig. S5B) are upregulated during the S phase, according to the KEGG enrichment. By contrast, the G2 phase is associated with higher activation of the glycolysis/gluconeogenesis and other carbohydrate-related pathways as well as sulfur metabolism. Many of the differentially expressed genes in G2 are implicated in glycolysis/gluconeogenesis and sulfur metabolism. These include sulfide:quinone oxidoreductase (SiRe_RS13005), implicated in sulfur metabolism, and phosphoenolpyruvate carboxykinase (SiRe_RS02025), aldose 1-dehydrogenase (SiRe_RS11380) and gluconate dehydratase (SiRe_RS10395), which participate in the glycolysis/gluconeogenesis and the pentose phosphate pathways (Fig. 3C). Because many of the enzymes in the glycolysis/gluconeogenesis pathway are shared with the fructose, mannose, sucrose and starch metabolism, the latter pathways were also enriched in the G2 phase (Supplementary Fig. S5C). However, no differential gene expression of enzymes specific for each of these pathways was found during the S and G2 phases. Moreover, because *S. islandicus* lacks the essential phosphofructokinase needed to perform glycolysis through the canonical Embden-Meyerhof-Parnas Pathway^46^, it is likely that glucose or fructose molecules are metabolized via the pentose phosphate pathway to glyceraldehyde or glycerate and then introduced into the glycolytic pathway to be fully oxidized. Finally, the profound shifts in the metabolic landscape during the G2 phase are also supported by the largest density of specifically co-expressed metabolism-related genes in this phase. In particular we observed specific co-expression of genes implicated in energy production and conversion and the carbohydrate metabolism genes (Supplementary Fig. S5A).

#### Cell division, motility and adhesion

As explained above, mitosis (M), division (D) and the pre-replicative first gap (G1) phases occur in rapid succession, precluding us from obtaining populations enriched in these discrete phases. Nevertheless, some of the proteins are known to be markers for the M and D phases. In particular, chromosome segregation during the M phase is mediated by a pair of proteins, SegA and SegB^47^, while cell division in the D phase is driven by the ESCRT-based machinery^27,48^. Consistently, during M-G1, we observed strong upregulation of genes encoding the pair of genome segregation proteins (Fig. 3A, Supplementary table S1) and ESCRT-based cell division machinery (including CdvA, ESCRT-III, ESCRT-III-1, ESCRT-III-2 and Vps4).

As in other members of the Sulfolobales^49–53^, motility and adhesion in *S. islandicus* are mediated by two evolutionarily related but functionally distinct extracellular filaments, the archaeal flagellum (or archaellum) and adhesive pili, respectively. Both filaments are composed of pilins/archaellins related to bacterial type IV pilins^54–56^ that are secreted through a membrane pore with the help of a cognate ATPase motors^57,58^. Live cell imaging of *S. acidocaldarius* cells revealed changes in the cell adhesion and motility around the time of division, suggesting the existence of coordination between these processes^59^. In particular, ∼80% of the observed cells underwent a transient loss of adhesion immediately prior or during cell division and >40% of newborn daughter cells rapidly moved away from the site of division^59^. Our data are fully consistent with these findings. Differential gene expression analysis showed that the pore and ATPase that secrete the archaellins are upregulated during the M-G1 and S phases (Fig. 4D). By contrast, the pore and ATPase responsible for the export of the adhesive pilins are both upregulated during the G2 phase (Fig. 4D). These patterns suggest that adhesive pili, present during the G2 phase, would be replaced by flagella following cell division. Interestingly, during the S phase, adhesive pilins and the ATPase of the archaellum were inversely co-expressed with an FKBP family peptidyl-prolyl cis-trans isomerase chaperone (SiRe_RS06340), suggesting the involvement of the latter in the switch between swimming motility and adhesion (Fig. 4E).

#### Defense systems

Our data shows a strong upregulation of various defense related genes during the S and G2 phases (Fig. 3A). Defense against viruses and mobile genetic elements in *S. islandicus* REY15A is primarily mediated by the CRISPR (clustered regularly interspaced short palindromic repeats) system^60^ (see Supplementary text for description of the *S. islandicus* CRISPR systems). In addition to the more extensively studied CRISPR-Cas systems, *S. islandicus* REY15A encodes two recently predicted, but functionally uncharacterized defense systems, namely, Hma^61^, composed of a helicase (SiRe_RS03020), methyltransferase (SiRe_RS03035) and ATPase (SiRe_RS03030), and the Methylation Associated Defense System (MADS) (SiRe_RS00305 and SiRe_RS00315)^62^ (Fig. 5A).

**Figure 5.**
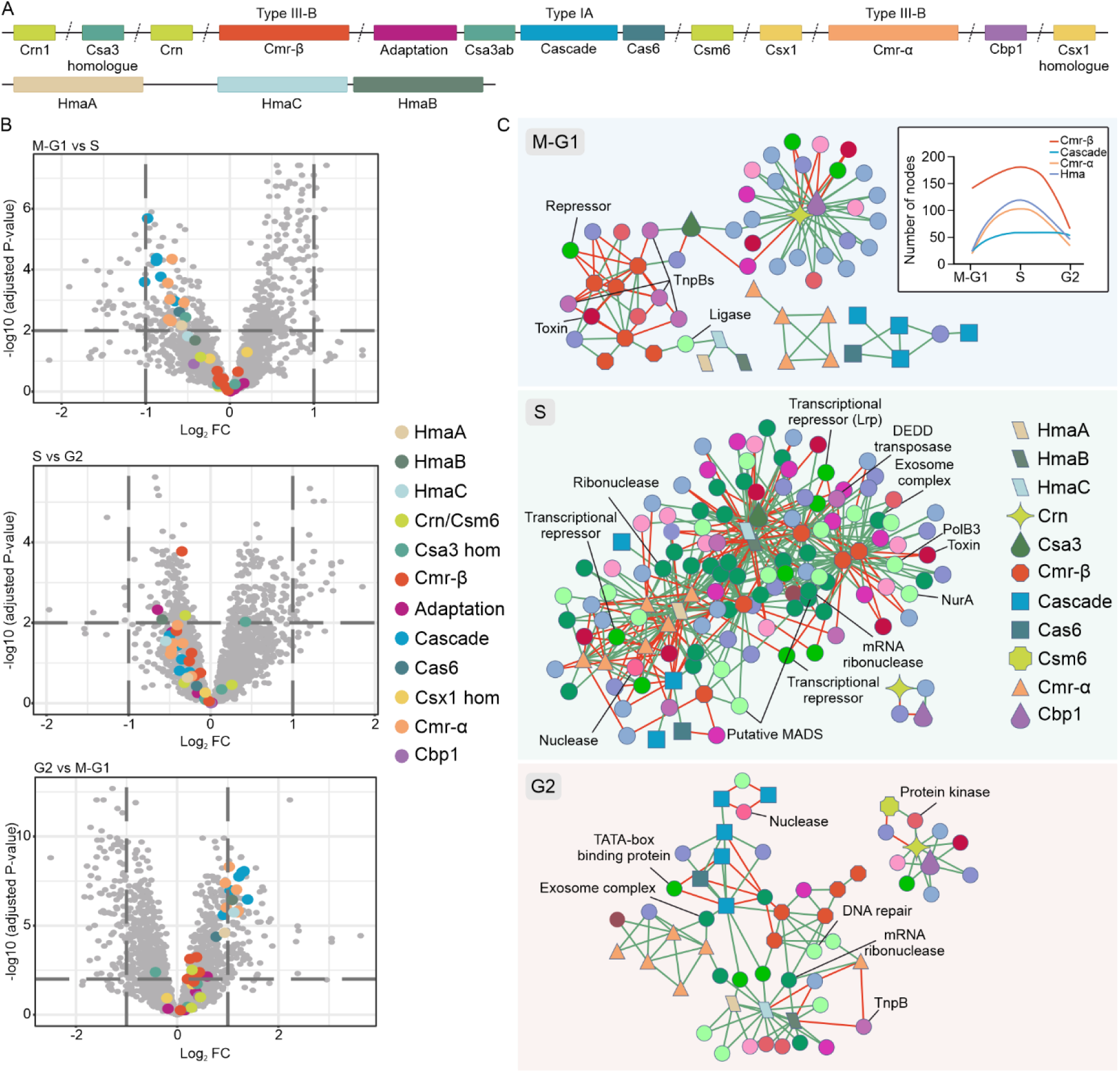
Defense systems are expressed cyclically. **A.** Organization of the different defense cassettes in the *S. islandicus* REY15A genome. CRISPR-Cas and Hma loci of REY15A are shown with different color-coded boxes. A diagonal dotted dash indicates a gap between the genes or cassettes. **B.** Expression of the different defense related genes and cassettes throughout the cell cycle. Pair-wise comparisons between phases are represented in three volcano plots. The y-axis represents the adjusted *p* value in logarithmic scale in base 10. The x-axis represents the fold change (FC) between the two phases in logarithmic scale in base 2. The horizontal lines mark the threshold for significance, i.e., a *p* value of <1 (-log_10_(0.01) = 2). The vertical lines mark the threshold for strong differential expression, i.e., FC of 2 (log_2_(±2) = ±1). Defense genes and cassettes are highlighted and color-coded. **C.** Gene co-expression subnetworks of the defense components. The subnetworks are extracted from the full GCNs by maintaining only those nodes that co-express with at least two defense-related (CRISPR-Cas or *hma*) genes. Nodes are colored by arCOG category following the same color code as in Fig. 2B, while edges are colored green if the co-expression is positive and red if it is negative. **C inset.** Size of the co-expression subnetwork of the CRISPR-Cas interference cassettes and the Hma cassette throughout the cell cycle. Line plot representing the number of nodes of the subnetwork of all genes co-expressing with the Cmr-β, the Cascade, the Cmr-α and the Hma cassettes at each of the phases of the cell cycle.

The differential gene expression analysis showed that different components of the Cascade complex, Cmr-α, and Cas6, which is responsible for the processing of the CRISPR RNA, were upregulated during the S phase, along with the Hma genes (Fig 5B). All the aforementioned genes are also active during G2, when they are joined by the upregulated Cmr-β genes. Such pattern of expression suggests activation of the defense and surveillance systems during S and G2, albeit with slight variation between the different CRISPR types and modules. Curiously, concomitant with their downregulation during the M-G1 and subsequent de-repression during the S phase, networks of co-expression of CRISPR related genes showed an inverse correlation with transcriptional repressors during these two phases (two Lrp family transcriptional regulators, SiRe_RS02835 and SiRe_RS07120, during M-G1 and S, respectively; and two ArsR family transcriptional regulators, SiRe_RS09190 and SiRe_RS09585, during S phase) (Fig. 5C). In addition, a PIN-domain ribonuclease toxin (SiRe_RS03095) and several transposases (IS*110* [SiRe_RS00565] and IS*5* [SiRe_RS04210] family transposases and TnpB family nucleases [SiRe_RS04095, SiRe_RS05390, SiRe_RS06735]) also showed similar co-expression, indicative of a possible control by the defense system. By contrast, positive correlation between the expression of Cas nucleases with diverse cellular nucleases, including mRNA ribonuclease (SiRe_RS03015), Rrp4 cap of the RNA-degrading exosome complex^63^ (SiRe_RS06470) and TatD-family nuclease (SiRe_RS05420), and proteins implicated in DNA repair, such as Mre11 (SiRe_RS00310), PolB3 (SiRe_RS09745) and NreA (SiRe_RS06100) suggests the presence of yet to be discovered cellular partners of the CRISPR-Cas systems. Some of these partner proteins could be, for instance, activated by cyclic oligoadenylate (cOA) as in the case of Csx1 or play a role in preserving genome integrity upon CRISPR-Cas activation. During the S phase, which contains the most coordinated network of the CRISPR-Cas systems (Fig. 5C inset), all three components of the Hma system and genes of the putative MADS defense co-expressed with different CRISPR-Cas modules suggesting coordination between these distinct defense systems.

It is generally considered that defense systems are under tight regulatory control and activated only upon invasion of foreign mobile genetic elements, such as viruses or plasmids^64,65^. Our results suggest that this might not be entirely the case. Activation of some of the defense systems during the S phase could be triggered by the exposed DNA replication intermediates or more active proliferation of transposons. Alternatively, CRISPR systems may additionally function in non-defense contexts, for instance, during DNA repair, as previously hypothesized^66^. For instance, it has been recently demonstrated that in halophilic archaeon *Haloferax volcanii*, Cas3 protein, component of type I systems, facilitates rapid recovery from DNA damage^67^. Indeed, our co-expression networks show coordination of DNA repair and defense systems, suggesting a role of the defense systems in safeguarding the integrity of the genome.

### Phase-specific signature genes defined using machine learning

To validate the results of the differential gene expression and GCN analyses, and to identify the signature genes defining each phase, we applied a machine learning algorithm to the transcriptomics and flow cytometry data (Supplementary table S4; see Methods). The flow cytometry data demonstrates that each sample, although enriched in cells from the targeted phase, includes cells from all three phases (Supplementary Figure S1). Using non-negative least squares (nnls) optimization we corrected gene expression levels in each sample by excluding transcription signal possibly originated from cells in non-targeted phases. *S. islandicus* genes were clustered based on corrected expression levels into four different groups depending on whether they tended to be more expressed at one of the three phases studied (groups 1-3) or display similar expression throughout the cell cycle (group 4) (Supplementary table S4). This allowed identification of gene sets showing phase-specific expression patterns. In the next step, to gain a clearer view on the processes taking place during each of the phases, we filtered out all the poorly annotated genes assigned to arCOG categories R or S (Supplementary table S4). The picture which emerged from this analysis was fully consistent with and complementary to the conclusions drawn from the manual analysis of the differential gene expression patterns and GCNs described in the previous sections (Fig. 6A). In particular, the signature genes specific of the M-G1 phase included those for genome segregation, cell division, DNA replication as well as nucleotide and lipid metabolisms; S-specific genes set included genes for diverse chromatin proteins and translation related genes as well as some of the carbon metabolism genes; G2 phase was characterized by genes for carbohydrate and sulfur metabolism. Notably, some of the signature genes identified by machine learning approach, did not stand out in the differential gene expression analysis. For instance, the signature gene set of the M-G1 phase includes the β-subunit of the proteasome and the proteasome-activating nucleotidase, a regulatory subunit that drives the conformational changes during the proteasome functional cycle^68^. It has been previously shown that in the presence of a proteasomal inhibitor, the ESCRT-III rings cannot be disassembled, resulting in cell division arrest in *S. acidocaldarius*^27^ and the same effect was observed in *S. islandicus*. These experimental results are consistent with the proteasomal genes being selected as the M-G1 signature genes. Thus, the three phase-specific gene sets defined using the machine learning approach appear to adequately represent the gene expression patterns along the *S. islandicus* cell cycle and can be used in subsequent analyses to assess the state of cells under different experimental conditions. Importantly, this analysis indicates that many processes in *S. islandicus* are coordinated with the cell cycle, being expressed during particular phases, resembling, at least qualitatively, the transcriptional landscape of the cell cycle in eukaryotes.

**Figure 6.**
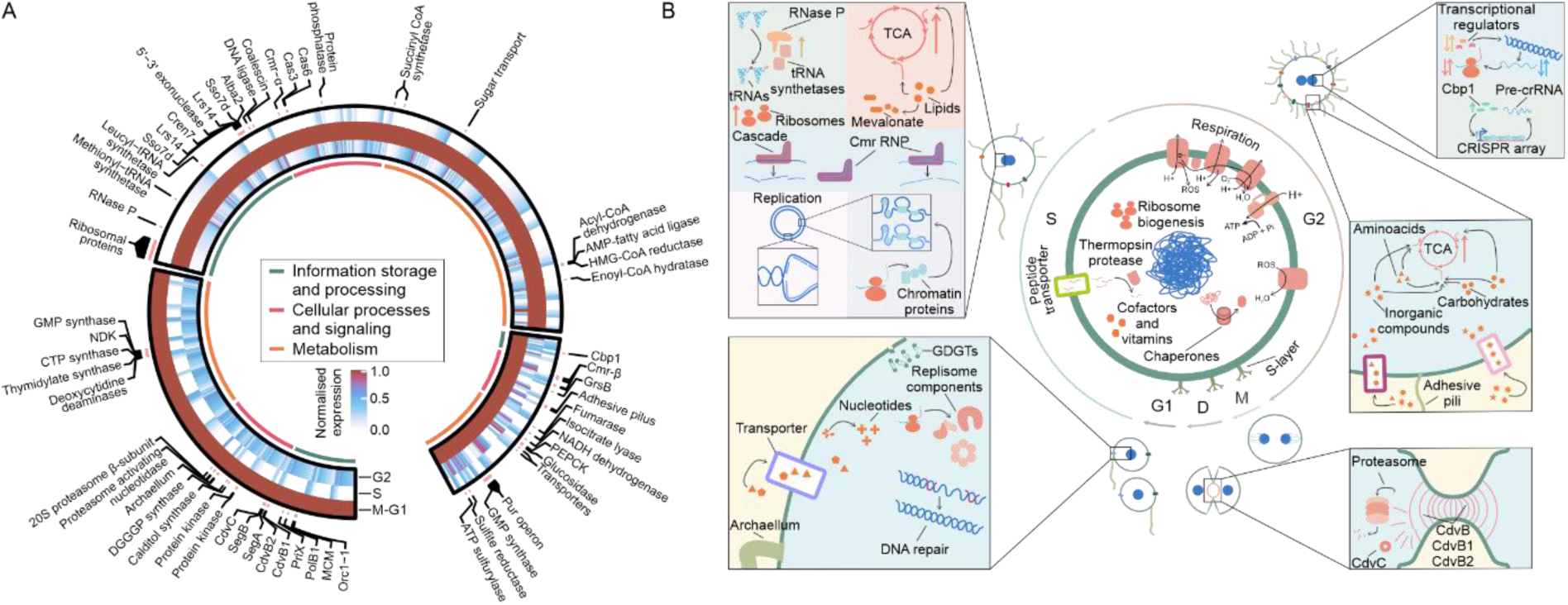
Cell cycle of *S. islandicus*. **A.** Circular heatmap of the signature genes identified by a machine learning approach. The predicted expression estimated by machine learning was normalized in between phases and genes were clustered based on k-means into four different groups depending on whether they tended to be more highly expressed at one of the three phases studied (groups 1-3) or displayed uniform expression throughout the cell cycle (group 4). The clusters of genes were represented in a circular heatmap with their normalized expression using the circlize R package. Genes are ordered by arCOG category and their arCOG class (information storage and processing, cellular processes and signaling or metabolism) is marked in the graph. In a clockwise order starting at 6 o’clock, genes clustering as peaking at M-G1 are shown first, then those peaking at S and finally those that do at G2. Genes of interest are marked with their annotations next to the heatmap. **B.** Graphical summary of the changes occurring during the cell cycle. The core and housekeeping processes which are maintained throughout the cycle are shown in the center of the graphic, whereas the phase-specific processes are depicted at the periphery. The presumed morphological states at each of the cell cycle phases are also depicted.

Notably, the identified signature genes were significantly (*p* value < 0.01) more conserved compared to the rest of the genes. Namely, 89.9% and 51.6% of the signature genes were conserved across Thermoproteota and three other archaeal phyla, respectively, whereas the non-signature genes displayed lower conservation in the corresponding lineages (78.6% and 44.5%, respectively). The higher conservation of the phase-specific genes suggests that the regulation and the overall structure of the *Saccharolobus* cell cycle is conserved in archaeal lineages beyond the order Sulfolobales. Finally, we assessed whether the expression of the *S. islandicus* phase-specific genes also follows a cell cycle dependent transcription pattern in eukaryotes. To this end, we compared the cell cycle phase affiliation of genes that are homologous between *S. islandicus* and eukaryotes represented in the Cyclebase database^69^ (see Methods). The budding yeast *Saccharomyces cerevisiae* shared the largest number (n=403) of homologs with *S. islandicus* (Table 1, Supplementary table S5). Of the 86 *S. islandicus* signature genes specific to the M-G1 genes, 67 homologous genes displayed peak expression during M or G1 phase in *S. cerevisiae*. Of the 57 G2 signature genes of *S. islandicus*, 37 were also expressed during G2 in the budding yeast. The S-specific genes, the largest category with 260 genes, 28.8% (n=75) of which encode ribosomal proteins and other translation or protein folding related proteins (tRNA synthetases, tRNA ligases, thermosome subunits, etc.), displayed less congruence in the temporal expression, with only 57 genes displaying peak expression during the S phase in both organisms (Table 1, Supplementary table S5). Thus, despite certain differences in the timing of expression of certain functions, the overall structuring of the transcriptional landscape follows a defined program in both *S. islandicus* and eukaryotes.

**Table 1.** Percentage of similarity between expression peaks of *S. cerevisiae* genes homologous to *S. islandicus* signature phase-specific genes.

| Cell cycle phase | Number of phase-specific *S. islandicus* genes with homologs in *S. cerevisiae* | % similarity (relaxed rules) | % similarity (strict rules) |
| --- | --- | --- | --- |
| M-G1 | 86 | 77.9 | 64.0 |
| S | 260 | 21.9 | 15.8 |
| G2 | 57 | 64.9 | 57.9 |

## CONCLUDING REMARKS: PARALLELS TO THE EUKARYOTIC CELL CYCLE

Our understanding of gene expression during the cell cycle in prokaryotes in general, and in archaea in particular is rather limited. In bacteria, gene expression appears to be primarily defined by the movement of the replication fork^70–73^. Our data suggest that this is not the case in *S. islandicus* or other Sulfolobales, where the expression program appears to be more akin to that of eukaryotes. In addition to the previously demonstrated cyclic expression of Sulfolobales genes related to cell division and DNA replication^25,28,31,48,74,75^, our results indicate that many other key cellular processes in *S. islandicus*, including defense systems, cell motility and adhesion apparatuses as well as diverse metabolic pathways, are coordinated with the progression of the cell cycle (Fig. 6B). Some of these processes are also expressed cyclically in eukaryotes^1,7,8,76^.

Despite the disparity in the duration of the G1 phase in eukaryotes^8^ and Sulfolobales, the overall ‘logic’ appears to be the same, that is, to prepare for genome replication. Indeed, our data suggests a transcriptional activation of DNA repair pathways and expression of the major replisome components, including the replicative DNA polymerase, MCM helicase and two of the three origin recognition genes (Orc1-1 and WhiP), during the M-G1. Similarly, the expression of the MCM subunits, helicase loader Cdc6 and origin recognition subunits (Orc1-6) in G1 have been demonstrated in eukaryotes^77^. In eukaryotes, G1 is the pivotal moment when the cell undergoes its fate: replication, differentiation, or death^1^. Whether a similar checkpoint exists in *S. islandicus* remains to be investigated. In eukaryotes, the G1 phase is also associated with high biosynthetic activity, accompanied by the increase in cell size^1^. Although in *S. islandicus*, most of the metabolic pathways displayed reduced activity during the M-G1, lipid biosynthesis was significantly enhanced, suggesting cell growth.

DNA replication marks the start of the S phase. In eukaryotes, cyclin-dependent kinases initiate replication through an orchestrated process with different origins^1,77^. Similarly, in Sulfolobales, the S phase starts with the almost simultaneous firing of two of the three origins of replication triggered by Orc1-1 and WhiP^78,79^, even though their transcripts are produced during the M-G1 phase (see above), consistent with the previous observations^74^. In eukaryotes, the DNA synthesis during the S phase is coordinated with the production of histones, facilitating the assembly of the newly replicated DNA into chromatin^80^. Remarkably, our data suggest that DNA replication and chromatinization are coupled in *S. islandicus*. We found upregulation of the dominant chromatin-associated proteins, including Cren7, Sul7d and Lrs14, during the S phase^44^. However, unlike in eukaryotes, where apart from histones, protein synthesis appears to be generally low^1^, we observed an upregulation of diverse translation-related genes, including those encoding the non-universally conserved core ribosomal proteins, tRNA synthetases, tRNAs and RNase P. Consistently, many genes encoding for ribosomal proteins and tRNA synthetases were found as signature genes of the S phase by machine learning approach. Consistent with the low translation during the S phase in eukaryotes, only 10 of the 75 translation related genes specific to the S phase in *S. islandicus* showed peak expression at the same phase in *S. cerevisiae*. Instead, most of the *S. cerevisiae* homologs showed peak expression during the M or G2 phases (Supplementary table S5). These differences may result from ultrastructural differences between eukaryotic and archaeal cells. In particular, presence of a nucleus in eukaryotes effectively uncouples transcription and translation, whereas in archaea, the two processes appear to be coupled^81,82^. Although protein levels were not measured in this study, in the future, it will be interesting to perform a cell cycle-resolved proteomics analysis and assess the correspondence between transcriptomics and proteomics.

One of the pronounced differences between the cell cycles of *S. islandicus* and eukaryotes concerns the G1 and G2 phase. The duration of the two phases appears to be inverted in the two models. In eukaryotes, G1 is one of the longest and G2 is one of the shortest phases^8^, whereas the opposite is true for Sulfolobales. In eukaryotes, G1 is the major growth phase, characterized by higher metabolic activity than the other phases^1^, whereas in *S. islandicus*, many metabolic pathways are most active during the S and G2 phases. By contrast, the G2 phase in eukaryotes is characterized by active lipid biosynthesis and DNA repair^1^, whereas these processes appear to be activated during M-G1 in *S. islandicus*. Nevertheless, despite these differences, the general logic of the coordination of metabolism, DNA repair, and lipid biosynthesis during one of the two G phases is shared. More importantly, most of the *S. islandicus* M-G1 and G2 signature genes that have homologs in *S. cerevisiae* showed their peak expression during the same phases in both organisms (Table 1). Thus, although the G1 and G2 phases in *S. islandicus* are compressed and elongated, respectively, compared to most eukaryotes, in terms of the executed cellular processes, the two phases appear to be equivalent in this archaeon and eukaryotes. Thus, it is tempting to speculate that the blueprint of the eukaryotic cell cycle has originated in archaea and has been maintained, albeit with notable modifications and regulatory adjustments, throughout the eukaryotic evolution.

Collectively, our results illuminate the complexity of the transcriptional landscape in an archaeal model system. Notably, the overall program of the cell cycle appears to be conserved throughout the class Thermoproteia^29^, suggesting that our findings could be extrapolated to other members of this archaeal lineage. In this context, the signature genes characteristic of different cell cycle phases could be particularly useful for future comparative studies.

## METHODS

### Strains and growth conditions

*Saccharolobus islandicus* strain REY15A was grown aerobically at 76°C with shaking in 25 ml of MTSV medium containing mineral salts (M), 0.2% (wt/vol) tryptone (T), 0.2% (wt/vol) sucrose (S) and a mixed vitamin solution (V); the pH was adjusted to 3.5 with sulfuric acid, as described previously^83^.

### Synchronization of Saccharolobus islandicus

Cells were synchronized using acetic acid (final concentration: 6 mM) as previously described^84^. Briefly, cells were cultured in 25 ml of MTSV media until OD_600_ reached 0.2, then cultures were synchronized by addition of acetic acid to arrest the cells at the end of the G2 phase. Following the incubation for 6h, the cells were pelleted at 2900xg for 15 minutes, washed with 0.7% sucrose to remove the acetic acid and resuspended in warm acetic-acid free media. Once synchronized, the cells were grown in the MTSV medium as described above and the progression of the cell cycle was followed by flow cytometry. Briefly, cells were fixed with 70% ethanol at +4°C and washed once with PBS. Fixed cells were stained with 40 μg/ml PI (Invitrogen) in staining buffer (100 mM Tris pH: 7.4, 0.5 mM NaCl, 1 mM CaCl_2_, 0.5 mM MgCl_2_, 0.1% Nonidet p-40) and their DNA content analyzed using CYTOflex (Beckman-Coulter).

### RNA extraction and sequencing

Cells were plated on solid media and 15 single colonies were inoculated in liquid media. The total RNA extracted from the 15 clonal cultures, which were considered as biological replicates, at three different time-points after synchronization: 2h30, 4h and 6h (Figure 1B). The time points were selected based on the results of the flow cytometry analysis which showed that samples at these three time points are enriched in cells in the M-G1, S and G2 phases, respectively (Figure 1C). Total RNA was extracted using TRI Reagent (SIGMA-Aldrich), following the manufacturer’s protocol, and treated with DNase (TURBO DNA-free kit; Invitrogen) following the manufacturer’s instructions. DNase treated samples were further purified with RNeasy Mini Kit (Qiagen). RNA samples were quantified using a Qubit Fluorometer (Thermo Fischer Scientific) and assessed for quality with a BioAnalyzer (Agilent). Libraries were prepared using the Illumina® Stranded Total RNA Prep, Ligation with Illumina® Ribo-Zero Plus kit, with custom ribodepletion. Following PCR amplification, all samples underwent two rounds of purification with AMPure beads (Beckman Coulter) to remove small fragments. Libraries were subsequently validated using both the Qubit Fluorometer and the Fragment Analyzer (Agilent). Sequencing was performed on an Illumina NextSeq 2000 sequencer with a P3 50-cycle kit and a target of 30 to 40 million reads per sample. The reads obtained were mapped to the reference *S. islandicus* REY15A genome (RefSeq accession number: NC_017276) with Bowtie-2^85^ using default parameters.

### Differential gene expression analysis

The amount of reads per gene was counted with featureCounts^86^ with default parameters. Data was then processed using R and the EdgeR library^87^. Read counts per gene were standardized to counts per million and filtered to eliminate all of the genes which did not have at least one read per million in all 15 samples. Finally, reads were normalized using the TMM method, which assumes that the majority of genes is not differentially expressed^88^. Logarithmic fold change in base two (log_2_FC) was calculated by subtracting the logarithmic average of two groups, e.g., the log_2_FC of M-G1 vs S is calculated by subtracting the logarithmic average of S to the logarithmic average of M-G1. The data was fitted to a linear model to calculate statistical significance using the limma package and *p* values were adjusted using the Benjamini-Hochberg method. Genes were considered up- or down-regulated if log_2_FC was at least ±0.5 (FC ±1) and the adjusted *p* value was <0.01, genes with a log_2_FC of at least ±1 (FC ±2) were considered to be strongly up- or down-regulated.

### KEGG enrichment

Information on the different metabolic pathways was extracted from the Kyoto Encyclopedia of Genes and Genomes (KEGG)^89^. KEGG enrichment *p* values were calculated by taking into account the adjusted *p* values of the genes in each pathway and comparing them with the genes outside the pathway by applying a Wilcoxon test. Significance threshold was set at 0.05. Gene ratio was calculated as the number of genes for which RNA sequencing data was available relative to the number of genes annotated in each pathway. Density plots were generated using the ggplot2 package in R with the transcriptomic data of the genes in each pathway.

### Gene annotation

Genes were annotated and classified using the Clusters of Orthologous Genes (arCOG) framework^36^, where each gene is assigned a code indexed to a specific category according to its orthologs in other archaea. Essentiality information was extracted from the previous study on the closely related *S. islandicus* strain M.16.4^37^. Information on compartmentalization of the genome was extracted from data obtained previously^35^. Statistical significance in the distribution of essential genes and compartments in the core network was calculated by performing a Fischer’s exact text comparing the distribution of two groups. For proteins of interest, the arCOG annotations were supplemented with the results from profile-profile comparisons using HHpred.

### Codon usage

Codon usage for each gene was calculated using the coRdon library in R (Elek A, Kuzman M, Vlahovicek K (2023). coRdon: Codon Usage Analysis and Prediction of Gene Expressivity. doi:10.18129/B9.bioc.coRdon, R package v1.20.0, https://bioconductor.org/packages/coRdon) using the coding sequences for REY15A extracted from GenBank (accession number: NC_017276). Average codon usage for the full genome was calculated using the Sequence Manipulation Suite^90^ using the bacterial (11) genetic code. Information on codon usage is provided in Supplementary table S3.

### Data visualization

Data was represented in the form of volcano plots, violin plots, box plots or dot plots using the ggplot2 package in R. Ggplot2 was used to generate and plot the regression model of the expression dependent on chromosome position. The method used was a generalized additive model.

### Gene coexpression networks

Read count matrices quantifying the gene expression in M-G1, S and G2 were normalized using the VST transformation from the DESeq2 R package. The normalized matrices were subsampled to generate the highest possible number of inferences of GCNs and to produce multiple gene co-expression networks (GCNs) for each phase of the cell-cycle; specifically, within 15 transcriptomes, 5 transcriptomes can be randomly sampled to generate 20 replicates with no more than 2 shared transcriptomes between any pair of replicates. For each cell cycle phase, this protocol returned 20 subsamples, from which 20 GCNs were built. In each GCN, nodes correspond to sequences assigned to genes, connected by edges. To build these GCN, we used the normalized count matrix as input for package WGCNA and the function core to compute the Pearson correlation coefficients (PCC). A high PCC threshold for edge inclusion in a GCN was set at 0.8 as a first step to avoid spurious correlations: the edges in each final filtered GCN were thus weighted by either strong positive (PCC > 0.8) or negative (PCC < -0.8) PCC values. Then, for each phase of the cell cycle, consensus GCNs were constructed using a majority rule that only retained high correlation edges present in more than 80% of the replicate GCNs. This strategy identified strongly correlated gene co-expression very commonly observed in a cell cycle phase and robust to sampling effects, because these co-expressions are observed almost irrespectively of what transcriptome samples were used to describe each cell cycle phase.

### Classification of genes by conservation level

To estimate the conservation of *S. islandicus* genes among archaeal groups, a reference archaeal protein sequence dataset was assembled from 292 complete representative genome assemblies out of all archaeal genomic assemblies recorded in the RefSeq database^91^ and all proteins of *S. islandicus* REY15A. Diamond^92^ was used to perform an all-against-all comparison (with parameters -e 1e-5 -k 1000) using this reference protein sequence dataset. Gene families were computed from the results of the Diamond search using a sequence similarity network, built by filtering results (using standard thresholding parameters: minimum % identity = 30%; minimum mutual sequence coverage = 80%), using the cleanblast and familydetector scripts from the MultiTwin tool^93^. The resulting gene families were mapped on a reference archaeal phylogeny^94^ using the ete3 Python package, and further categorized by conservation level, based on the phylogenetic distribution of their members. Differences in conservation levels between genes in the core network compared to non-core genes was tested by performing a Chi-square test comparing the conservation distribution of the non-core genome with the core genes.

### Identification of phase-specific signature genes

To estimate the signature genes for each phase, we assumed that each sample consisted of three subpopulations of cells: (i) M-G1, (ii) S or (iii) G2. Each subpopulation has a different proportion of cells: *p*_*M*−*G*1_ + *p*_*S*_ + *p*_*G*2_ = 100% ( *p*_*M*−*G*1_, *p*_*S*_, *p*_*G*2_ = percentage of cells in the M-G1, S and G2 subpopulations, respectively). Moreover, we assumed that each gene has a constant transcription level (number of reads) in all cells from one subpopulation (*t*_*M*−*G*1_, *t*_*S*_, *t*_*G*2_ = transcription levels of a gene in the subpopulations M-G1, S and G2). With these assumptions, the total transcription level of a gene (*T*) can be presented as a sum of the transcription levels from three subpopulations of cells: *T* = *p*_*M*−*G*1_ ∗ *t*_*M*−*G*1_ + *p*_*S*_ ∗ *t*_*S*_ + *p*_*G*2_ ∗ *t*_*G*2_. The total transcription level or number of reads of a gene (*T*) was calculated with featureCounts (see above). The percentage of cells in different subpopulations (*p*_*M*−*G*1_, *p*_*S*_, *p*_*G*2_) was obtained from the flow cytometry data (see above) of three representative biological replicates (Supplementary Fig. S2A; Supplementary table S4). From this data, for each gene we have nine linear equations (*T* = *p*_*M*−*G*1_ ∗ *t*_*M*−*G*1_ + *p*_*S*_ ∗ *t*_*S*_ + *p*_*G*2_ ∗ *t*_*G*2_) with three unknown variables (*t*_*M*−*G*1_, *t*_*S*_, *t*_*G*2_). The unknown variables were predicted using the non-negative least square method in python (nnls function scipy version 1.11.4). As a result, for each gene we estimate *t*_*M*−*G*1_, *t*_*S*_, *t*_*G*2_, the transcription levels or number of reads in each phase. Once the reads were predicted, we excluded all the genes whose expression was estimated to be lower than 3 counts per million reads in all phases. The remaining genes were clustered based on K-means into four different groups depending on whether they were more expressed at one of the three studied phases (groups 1-3) or displayed a similar expression throughout the cell cycle (group 4). A second round of clustering was performed excluding all poorly annotated genes (arCOG categories R and S).

### Phylogenetic distribution of *S. islandicus* phase-specific genes

To determine whether *S. islandicus* phase-specific genes have homologs within bacterial and/or eukaryotic genomes, a reference proteic sequence dataset was built using all the reference proteomes from Uniprot^95^ for Bacteria (9285 proteomes) and Eukaryota (2625 proteomes). Diamond^92^ was used to perform a search (with parameters -e 1e-5 -k 10000) for homologs of 1530 proteins encoded by *S. islandicus* phase-specific genes against this reference protein sequence dataset. Depending on the phylogenetic diversity in homolog gene sets, *S. islandicus* phase-specific genes were classified as Prokaryotic if shared with bacteria only, Archaeoeukaryotic if shared with eukaryotes only or Mixed if shared with both groups.

### Comparison of the expression peaks for homologous phase-specific *S. islandicus* and eukaryotic genes

To compare the cell cycle-dependent gene expression patterns in archaea and eukaryotes, *S. islandicus* phase-specific genes eukaryotic homologs were identified in the CycleBase database^69^, which records the expression profile of periodically expressed genes during the eukaryotic cell cycle for 4 eukaryotic species (*S. cerevisiae*, *S. pombe*, *H. sapiens* and *A. thaliana*). A blastp search was performed (with parameters -evalue 1e-5 -word_size 5) using the proteins encoded by *S. islandicus* phase-specific genes as query against an eukaryotic protein sequence dataset combining the proteins from Uniprot reference proteomes for *S. cerevisiae* (UP000002311), *S. pombe* (UP000002485), *H. sapiens* (UP000005640) and *A. thaliana* (UP000006548). Since hits were mostly found within *S. cerevisiae* proteins, expression peak times were then compared between *S. islandicus* and *S. cerevisiae*; for each *S. islandicus* phase-specific signature gene, similarity to *S. cerevisiae* cell cycle transcriptomic data was decided when at least one of the three closest periodically expressed *S. cerevisiae* homologs (best blast hits) was maximally expressed at a corresponding phase, according to either relaxed (correspondences: archaeal M-G1 with eukaryotic G1, G1/S, G2/M and M phases; archaeal S with eukaryotic G1/S and S phases; archaeal G2 with eukaryotic G2 and G2/M phases) or strict (correspondences: archaeal M-G1 with eukaryotic G1 and M phases; archaeal S with eukaryotic S phase; archaeal G2 with eukaryotic G2 phase) phase correspondence rules.

## DATA AVAILABILITY

The raw reads generated in this study were deposited in European Nucleotide Archive under the accession number PRJEB75364.

## ACKNOWLEDGEMENTS

This work was supported by Agence Nationale de la Recherche grant ANR-23-CE13-022 to MK. The work in EB laboratory was supported by an ATM grant from the MNHN (ATM AAP 2023) and an Emergence grant from Sorbonne Université (S21JR31001—IP/S/V2 EMERG-ESPA). MGRV was supported by a stipend from the Pasteur-Paris University (PPU) International PhD Program. We also acknowledge the help of Pierre-Henri Commere and the Flow Cytometry platform at Institut Pasteur. The Biomics Platform, C2RT, Institut Pasteur, Paris, France, is supported by France Génomique (ANR-10-INBS-09) and IBISA.

## SUPPLEMENTARY INFORMATION

**Supplementary Figure S1.**
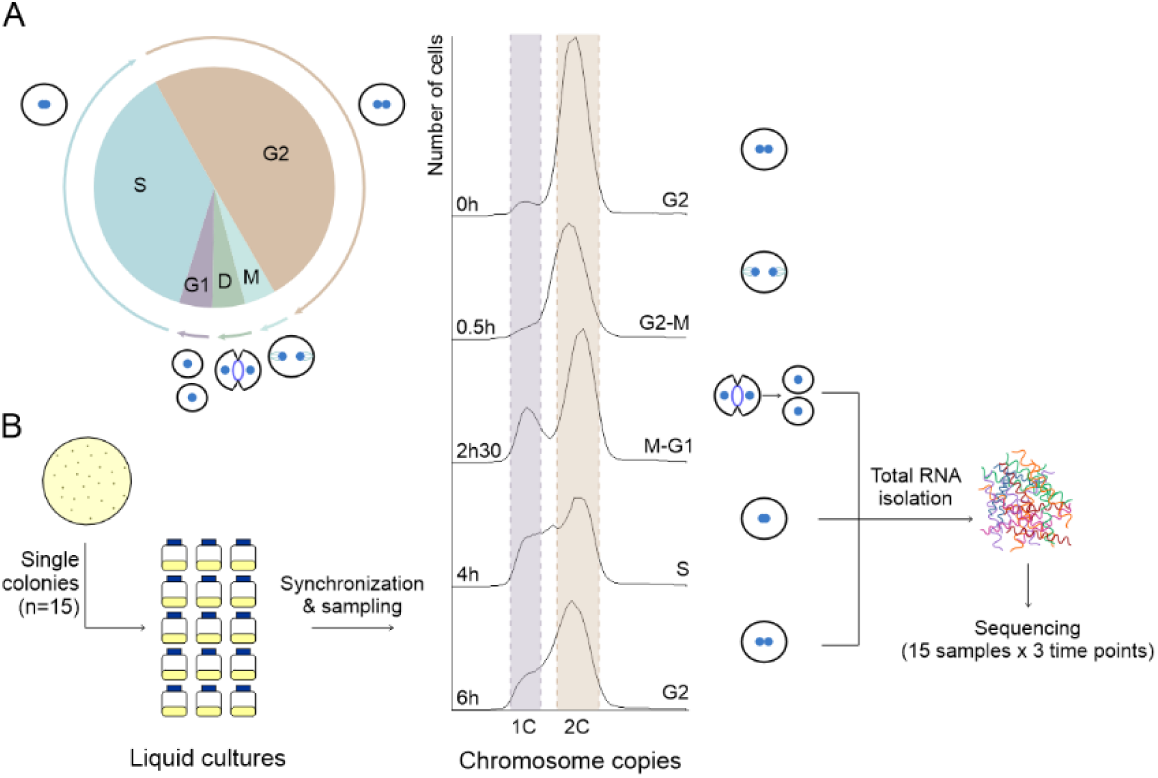
Schematic representation of the cell cycle and experimental workflow. **A.** The cell cycle of an exponentially growing *Saccharolobus* cell. Cell cycle phases occupy an area proportional to their typical duration. Morphological changes occurring throughout the cell cycle are depicted next to the corresponding phases. **B.** Experimental workflow for the transcriptomic analysis. Fifteen *S. islandicus* colonies were isolated and inoculated in liquid medium. The cultures were synchronized with acetic acid and following the cell cycle arrest release, samples were collected at the indicated time points during which the population was enriched in cells undergoing a particular cell cycle phase. The cell cycle phase at each sampled time point was determined by evaluating the genomic content of the cells using flow cytometry in three replicates (a representative profile is shown). The presumed morphological states characterizing different cell cycle phases are shown next to the corresponding flow cytometry profiles. The total RNA was extracted from each of the 15 cultures during the time points corresponding to the enrichment in cells undergoing the M-G1, S and G2 phases. The RNA was sequenced using Illumina platform.

**Supplementary Figure S2.**
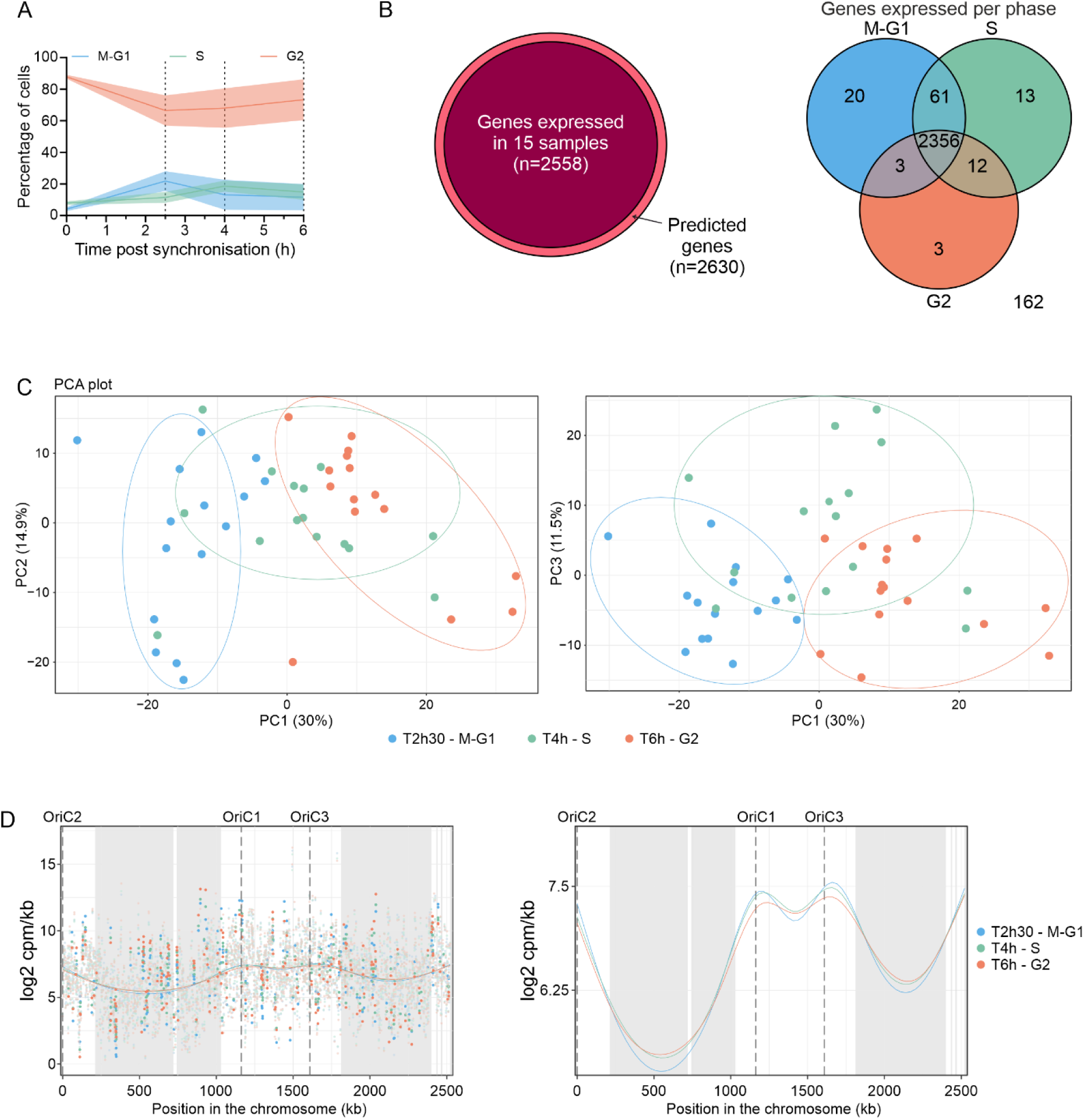
Overview of the differential expression analysis. **A.** Percentage of cells at each cell cycle phase during the sampling time frame. Line plot showing the percentage of cells progressing through each of the phases at each sampled time point. Phase was determined by evaluating the genomic content of the cells using flow cytometry: one chromosome copy corresponds to G1, two chromosome copies to G2 and cells in between, to S. Plot shows the average and standard deviation of three representative replicates. Dashed vertical lines mark the sampled time points (2h30, 4h and 6h). **B.** Venn diagrams showing the number of expressed genes. Left panel shows that 2558 of the 2630 predicted genes are expressed in at least 15 samples, these genes are included in the analysis. Right panel shows that of the 2630 predicted genes, 2356 are expressed in all phases in all 15 biological replicates, 112 are expressed only in one or two phases, while 162 are not expressed in any of the phases in all 15 biological replicates. **C.** Principal component analysis. Scatter plots show a graphical representation of the three principal components that explain most of the variance between the samples. Samples are colored by sampled time point. Each axis represents a principal component (PC) with the percentage of variance explained by that component shown in parentheses. Circles show the clustering of the different samples depending on the sampled time point. **D.** Scatter plots show the expression in counts per million per kilobase in logarithmic scale in base 2 of each gene positioned along the *S. islandicus* REY15A chromosome. Plots show the average expression of the 15 replicates at each sampled time point, each time point is color coded. Strongly differentially expressed genes are highlighted. Information on compartmentalization of the genome was extracted from data obtained previously^35^. The B compartment is shown with a grey background, while the A compartment is shown in white. Horizontal dashed lines mark the position of the three origins of replication. A regression model generated by the generalized additive model is plotted for each time point. The line plot on the right shows a magnified version of the regression models.

**Supplementary Figure S3.**
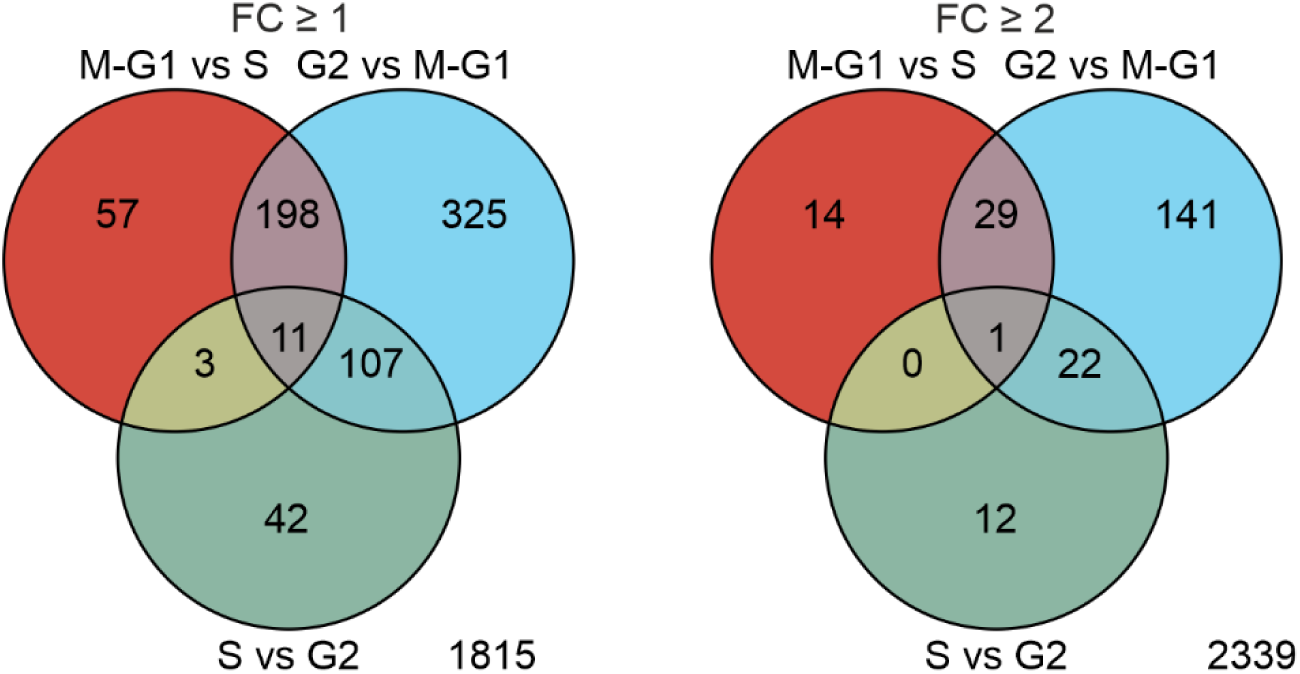
Overview of the pair-wise comparisons in the differential expression analysis. Venn diagrams show the genes differentially expressed (fold change [FC] ≥ 1, left) and strongly differentially expressed (FC ≥ 2, right) at each of the three pair-wise comparisons: (i) M-G1vsS, (ii) SvsG2 and (iii) G2vsM-G1. Briefly, from a total of 2558 analyzed genes, 743 were differentially expressed, 219 of those strongly.

**Supplementary Figure S4.**
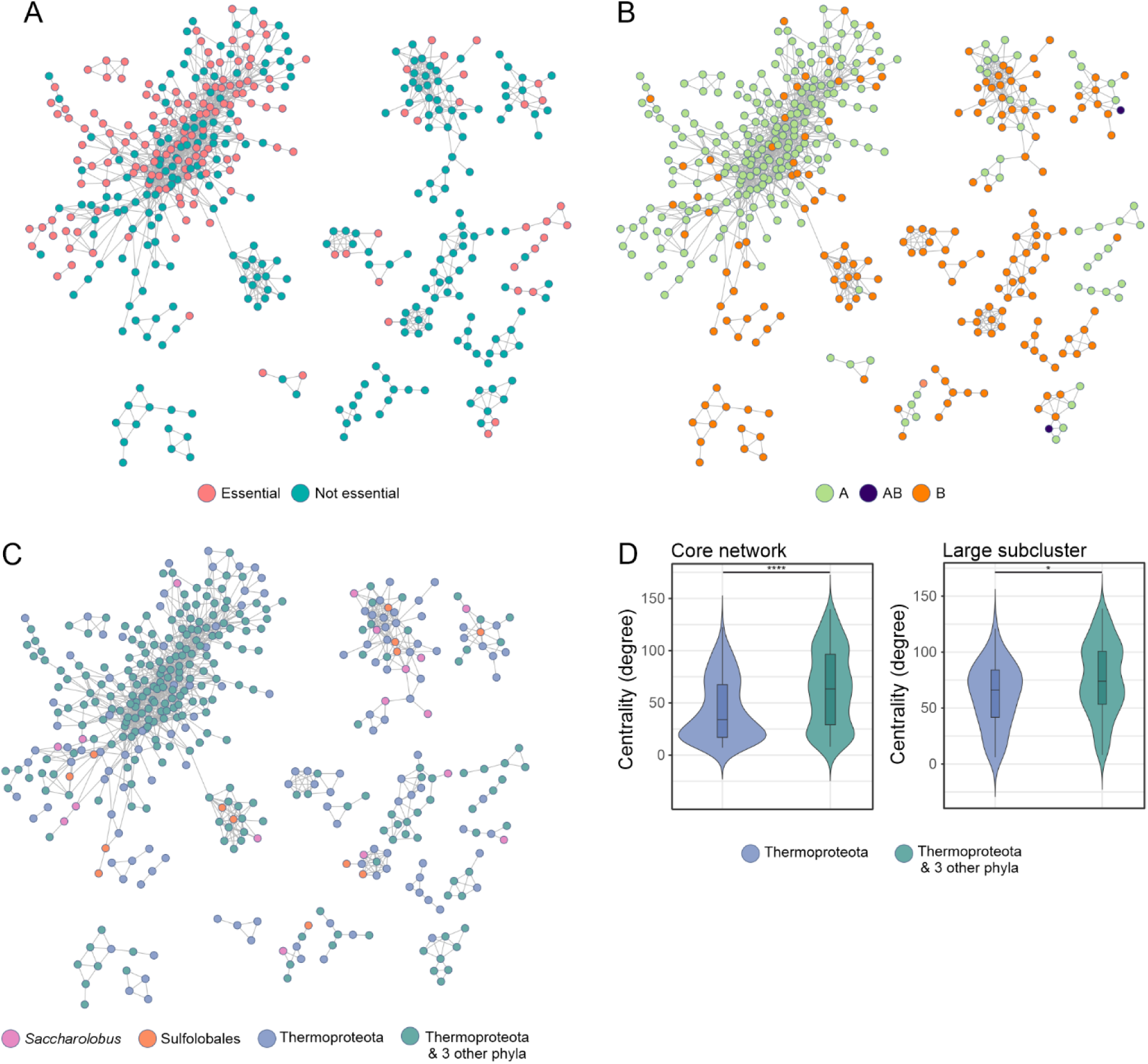
Topological analysis of the core network. **A.** Topological representation of the core network in which genes are colored according to their essentiality. Essentiality information was extracted from the previous study on the closely related *S. islandicus* strain M.16.4^37^. **B.** Topological representation of the core network in which genes are colored according to their compartment. Information on compartmentalization of the genome was extracted from data obtained previously^35^. **C.** Topological representation of the core network in which genes are colored according to their conservation. Genes were assigned to one of the four categories according to their conservation: (i) genes exclusive to the genus *Saccharolobus*, (ii) those restricted to the order Sulfolobales, (iii) genes conserved across the phylum Thermoproteota, and (iv) genes present in Thermoproteota and three other archaeal phyla, namely, Methanobacteriota, Halobacteriota and Thermoplasmatota. **D.** Centrality of the genes is dependent on their conservation. Violin plots show the degree of centrality of the different genes in the core network (top) and in the large subcluster of the network (bottom). Statistical significance of the difference in the average is calculated through a Wilcoxon test. * = *p* value <0.05, **** = *p* value <0.00005.

**Supplementary Figure S5.**
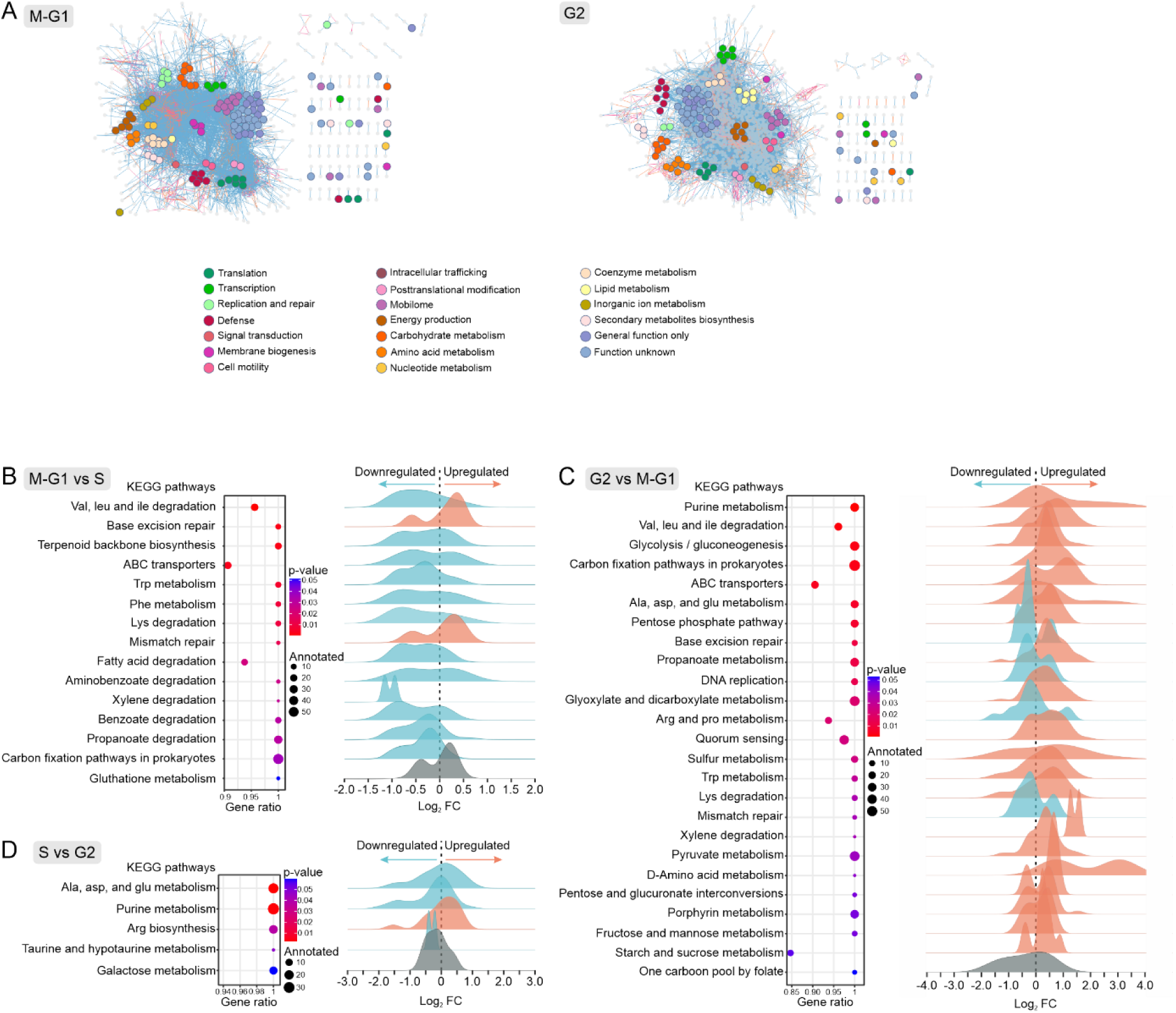
Cell cycle phase specific processes. **A.** Specific co-expression during the M-G1 and G2 phases. GCNs of the M-G1 (left) and G2 (right) phases with the specific nodes highlighted, grouped and colored by arCOG category. **B-D.** KEGG enrichment analysis of the three pair-wise comparisons (M-G1vsS, SvsG2 and G2vsM-G1). A representation summarizing the KEGG enrichment analysis is shown for each of the three pair-wise comparisons performed in the DGE analysis. Each graph shows the pathways that are significantly enriched (*p* value <0.5) in order of significance with the last pathway being the first non-significant. Dot plots show the gene ratio (number of genes analyzed in the dataset compared to the total genes annotated for the pathway) in the x-axis. The size of the dots is dependent on the number of annotated genes for each pathway and the color on the value of the *p* value. Density plots show the distribution of the fold change (FC) in all the genes at each pathway. The x-axis represents the FC in logarithmic scale in base 2. The dashed vertical line marks the 0 FC, with negative values meaning there is downregulation and positive values indicating upregulation. The plots are colored depending on whether the pathway is upregulated (orange), downregulated (blue) or not significant/undetermined (grey).

## Supplementary text

### Protein synthesis

Most genes implicated in translation, including those encoding for most of the ribosomal proteins, are expressed throughout the cell cycle and are part of the core network. Nevertheless, some components of the protein synthesis machinery, including some of the ribosomal proteins, especially those that are not part of the universally conserved ribosomal core^96,97^, displayed distinct expression patterns. In particular, non-core ribosomal proteins eL39 (SiRe_RS08375) and eS19 (SiRe_RS08365) were upregulated during the S phase (Supplementary table S2), whereas eL8 (SiRe_RS09700) and eS17 (SiRe_RS08705) were upregulated during the M-G1 and S phases, and eL20 (SiRe_RS08390) during the S and G2 phases. All the archaea-specific ribosomal proteins characterized to date^98,99^, namely, aL45 (SiRe_RS05075), aL46 (SiRe_RS05770), aL47 (SiRe_RS07130) and aS21 (SiRe_RS09330), were upregulated during the M-G1 and S phases, with the exception of aL47, which showed a similar expression pattern but the change was not statistically significant. The only core ribosomal protein differentially expressed during the cell cycle phases, uL29 (SiRe_RS06575), was downregulated during M-G1. These results suggest the maintenance of a stable ribosomal core to which different non-core components are added during particular cell cycle phases, possibly to finetune the efficiency and specificity of protein synthesis.

Notably, certain tRNA genes are strongly upregulated during the M-G1 and S phases (Fig 3A, Supplementary table S1). These include tRNA-Met^ATG^, tRNA-Trp^TGG^, tRNA-Asp^GAC^, tRNA-Ala^GCC^ and tRNA-Gln^CAG^ (Supplementary table S3). The upregulation of specific tRNAs suggests an additional layer in the regulation of protein synthesis. Several genes are indeed enriched in the codons recognized by the upregulated tRNAs and some are upregulated during the S phase, e.g., two genes encoding Sso7c4 homologs of the AbrB/MazE/SpoVT family DNA-binding proteins (SiRe_RS07595 and SiRe_RS09950). However, we could not recognize a consistent, unifying pattern which would rationalize the coordinated expression of these genes (Supplementary table S3). Furthermore, a glycyl-tRNA synthetase (SiRe_RS07915), a subunit of the RNase P (SiRe_RS06580) and the translation initiation factor 6 (SiRe_RS08385), which has been shown to exert translation control in eukaryotes^100^, were differentially expressed with their peak of expression being during the S phase (Supplementary table S2). Finally, the translation related genes are proportionally more prevalent in the S phase-specific GCN (Fig. 3B), further suggesting higher coordination and enrichment of protein synthesis during this phase.

### DNA repair

The M-G1 phase is also characterized by upregulation of the DNA repair genes, as is evident from the KEGG enrichment analysis (Supplementary Fig. S5B). Furthermore, the M-G1 GCN showed a higher proportion of specific co-expression of DNA repair-related genes (Supplementary Fig. S5A), suggesting that DNA repair takes place during the pre-replicative phase. In particular, endonuclease IV (SiRe_RS13470), an important player in base excision repair, is upregulated during M-G1, with uracyl-DNA glycosylase (SiRe_RS11945) and endonuclease III (SiRe_RS09590) showing less strong but similar trend. By contrast, mismatch specific nuclease, EndoMS/NucS^101,102^ (SiRe_RS00120), was not strongly upregulated during M-G1. These results suggest that only base excision repair is specifically active during the M-G1 phase, whereas the mismatch repair could be regulated by other specific cues. Given that most of the replisome components may also participate in the DNA repair processes, their expression during the M-G1 could signify a more active role in repair during M-G1, rather than being produced in advance and deployed only during the S phase.

### Amino acid and nucleotide metabolism

Many pathways related to the synthesis of amino acids are more active during the S and G2 phases (namely, the metabolism of alanine, aspartate, glutamate, isoleucine, leucine, lysine, phenylalanine, tryptophan and valine) (Fig. 3C, Supplementary Fig. S5B-D). Some of these pathways are also differentially activated between the S and G2 phases. For instance, the metabolism of alanine, aspartate and glutamate is more active during the G2 phase. By contrast, arginine biosynthesis is more active during the M-G1 and S phases (Supplementary Fig. S5C). In particular, the carbamoyl-phosphate synthase (SiRe_RS06955 and SiRe_RS06960) and the argininosuccinate synthase (SiRe_RS06970), which synthesize carbamoyl-phosphate and modify it inside the urea cycle into arginine, are upregulated during the M-G1 and S phases, whereas other enzymes of the pathway follow a similar but not significant trend.

Some of the key enzymes involved in nucleotide synthesis, such as the thymidylate synthase, exclusively implicated in DNA synthesis, or the CTP synthase, are specifically upregulated during M-G1 (Fig 3C). Furthermore, carbamoyl-phosphate, whose synthesis is enhanced during the M-G1 phase (see above), can also be a substrate for the synthesis of pyrimidines de novo. Somewhat unexpectedly, the synthesis of purines appears to be more active during the G2 (Fig. 3C, supplementary fig S5D-E). Enzymes encoded by the *pur* operon and functioning at the onset of the purine biosynthesis pathway are strongly upregulated during the G2 phase (Supplementary table S1). The lack of strict coordination between the biosynthesis of purines and pyrimidines might be linked to the fact that beside their role in the synthesis of nucleic acids, the pool of purines is also consumed for the energy storage in the form of ATP and GTP as well as synthesis of diverse coenzymes and signaling molecules, including FAD, NAD, cAMP, cOA, etc.

### S. islandicus CRISPR systems

*S. islandicus* REY15A carries three CRISPR-Cas effector modules: one type I-A (Cascade) and two type III-B Cmr modules, Cmr-α and Cmr-β^103^. The three effector modules, responsible for cleavage of the invading DNA or RNA, share the adaptation module, which is encoded next to the type I-A locus and consists of proteins Cas1, Cas2 and Cas4, required for acquisition of spacers from the invading foreign DNA and their insertion into the CRISPR arrays (Fig. 5A)^104^. The expression of the three CRISPR systems is tightly controlled and intricately regulated, but the underlying mechanisms are not fully understood. One of the central players in this regulation is cyclic oligoadenylate (cOA), second messenger synthesized by the type III-B systems in response to infection^105^. The cOA activates a range of ancillary defense proteins, such as Csx1 family nucleases, that nonspecifically cleave both viral and host mRNA^106,107^. In addition, cOA regulates the activity of the CARF domain-containing Csa3 family transcription factors (Fig. 5A)^108,109^. The levels of cOA return to normal after infection clearance thanks to Crn (CRISPR ring nuclease) family ring nucleases, which cleave cOA thereby deactivating the defense systems and preventing their cytotoxicity^110^. Finally, the efficient transcription of long CRISPR arrays is regulated by a CRISPR-specific transcription factor Cbp1^111,112^.

### Supplementary tables

**Supplementary Table S1. Differential gene expression analysis.** Dataset containing the results from the differential gene expression analysis. **Annotations**. Contains the annotation of all the predicted genes with their arCOG categories and information on compartmentalization, essentiality and conservation. **arCOG categories.** The indexed letter code for the different arCOG categories. **Expression Yes or No.** Expression of the predicted genes in the dataset. **GCN nodes-edges.** The number of nodes and edges of each category at each of the three GCNs. **DGE M-G1vsS; DGE SvsG2; DGE G2vsM-G1.** Results of the different pair-wise comparisons between phases with the fold change (FC) in logarithmic scale in base 2, the average expression and different statistics, including the adjusted *p* value. **Pseudogenes**. Pseudogenes that were excluded from the differential gene expression analysis.

**Supplementary Table S2. Core network.** Dataset containing all the information on the core network of *S. islandicus*. **Core network.** All the genes included in the core network with their different annotations, as well as their degree of centrality, whether they differentially express and information on the universally conserved ribosomal core. **Largest subcluster**. Genes included in the large subcluster of the core network. **Compartment; arCOGs; Level of conservation; Essentiality.** Information on the number of genes pertaining to the different compartments, arCOG categories, conservation levels and essentiality, respectively.

**Supplementary table S3. Codon usage.** Information on the codon usage in *S. islandicus*. **Full genome.** Codon usage in the full genome of *S. islandicus*. **Codon usage per gene.** Number of times each codon appears in each gene. **Codon usage per gene normalize.** Number of times each codon appears in each gene normalized by size of the gene. **GCC-Ala; CAG-Gln; GAC-Asp; TTG-Leu; ATG-Met; TGG-Trp.** The first ten genes with the highest prevalence of each of the analyzed codons along with their annotations.

**Supplementary table S4. Signature genes estimated through machine learning.** Prediction through machine learning of the expression of the different genes and clustering. **Percentage of cells per phase.** Number and percentages of cells that correspond to each of the three phases (M-G1, S and G2) in the three measured representative replicates. The number of cells was measured by flow cytometry and each phase assigned according to the genomic content (i.e. one copy of the chromosome is M-G1, two copies of the chromosome is G2 and between one and two copies is S). **Expression by phase log scale**. Predicted expression of each gene through a machine learning approach. Expression is shown in counts per million reads in logarithmic scale in base 2. Residuals indicate the error margin in counts per million reads. Maximum expression is the highest expression of the gene throughout the cell cycle. All of the genes whose expression does not surpass the minimum of 3 counts per million in logarithmic scale (8 counts per million) are excluded from further analysis. **Absolut expression normalize.** Predicted expression of each gene in counts per million with the ratio at each of the phases and their clustering by K-means.

**Supplementary table S5.** Comparison of the cell cycle phase affiliations of signature genes shared by *S. islandicus* and *S. cerevisiae*. Table is color coded to show when the homologs peaked at the same or similar phases in *S. islandicus* and *S. cerevisiae*.

